# Computation of the electroencephalogram (EEG) from network models of point neurons

**DOI:** 10.1101/2020.11.02.364802

**Authors:** Pablo Martínez-Cañada, Torbjørn V. Ness, Gaute T. Einevoll, Tommaso Fellin, Stefano Panzeri

**Author notes:** Corresponding author (S.P.).

## Abstract

The electroencephalogram (EEG) is one of the main tools for non-invasively studying brain function and dysfunction. To better interpret EEGs in terms of neural mechanisms, it is important to compare experimentally recorded EEGs with the output of neural network models. Most current neural network models use networks of simple point neurons. They capture important properties of cortical dynamics, and are numerically or analytically tractable. However, point neuron networks cannot directly generate an EEG, since EEGs are generated by spatially separated transmembrane currents. Here, we explored how to compute an accurate approximation of the EEG with a combination of quantities defined in point-neuron network models. We constructed several different candidate approximations (or proxies) of the EEG that can be computed from networks of leaky integrate-and-fire (LIF) point neurons, such as firing rates, membrane potentials, and specific combinations of synaptic currents. We then evaluated how well each proxy reconstructed a realistic ground-truth EEG obtained when the synaptic input currents of the LIF network were fed into a three-dimensional (3D) network model of multi-compartmental neurons with realistic cell morphologies. We found that a new class of proxies, based on an optimized linear combination of time-shifted AMPA and GABA currents, provided the most accurate estimate of the EEG over a wide range of network states of the LIF point-neuron network. The new linear proxies explained most of the variance (85-95%) of the ground-truth EEG for a wide range of cell morphologies, distributions of presynaptic inputs, and position of the recording electrode. Non-linear proxies, obtained using a convolutional neural network (CNN) to predict the EEG from synaptic currents, increased proxy performance by a further 2-8%. Our proxies can be used to easily calculate a biologically realistic EEG signal directly from point-neuron simulations and thereby allow a quantitative comparison between computational models and experimental EEG recordings.

**Author summary:** Networks of point neurons are widely used to model neural dynamics. Their output, however, cannot be directly compared to the electroencephalogram (EEG), which is one of the most used tools to non-invasively measure brain activity. To allow a direct integration between neural network theory and empirical EEG data, here we derived a new mathematical expression, termed EEG proxy, which estimates with high accuracy the EEG based simply on the variables available from simulations of point-neuron network models. To compare and validate these EEG proxies, we computed a realistic ground-truth EEG produced by a network of simulated neurons with realistic 3D morphologies that receive the same spikes of the simpler network of point neurons. The new obtained EEG proxies outperformed previous approaches and worked well under a wide range of simulated configurations of cell morphologies, distribution of presynaptic inputs, and position of the recording electrode. The new proxies approximated well both EEG spectra and EEG evoked potentials. Our work provides important mathematical tools that allow a better interpretation of experimentally measured EEGs in terms of neural models of brain function.

## Introduction

Electroencephalography is a powerful and widely used technique for non-invasively measuring neural activity, with important applications both in scientific research and in the clinic [1]. Electroencephalography has played a key role in the study of how both neural oscillations and stimulus-evoked activity relate to sensation, perception, cognitive and motor functions [2–4]. The electroencephalogram (EEG), like its intracranial counterpart, the local field potential (LFP), originates from the aggregation of all the electric fields generated by transmembrane currents across the surfaces of all neurons sufficiently close to the electrode [5–8]. The physics of how electromagnetic fields are generated from transmembrane currents are well understood, and mathematically described by forward models [6]. Yet, how to interpret changes in EEG across experimental conditions or diagnostic categories in terms of underlying neural processes remains challenging [1].

One way to better understand the EEG in terms of neural circuit mechanisms and to link theoretical models of brain functions to empirical EEG recordings is to compare EEG data with quantitative predictions obtained from network models. Network models of recurrently connected leaky-integrate-and-fire (LIF) point neurons are a current major tool in modelling brain function [9–11]. These models reduce the morphology of neurons to a single point in space and describe the neuron dynamics by a tractable set of coupled differential equations. These models are sufficiently simple to be understood thoroughly, either with simulations that are relatively light to implement, or by analytical approaches [12, 13]. Despite their simplicity, they generate a wide range of network states and dynamics that resemble those observed in cortical recordings. They have been employed to satisfactorily explain a broad spectrum of different cortical mechanisms and cortical functions, such as sensory information coding [14, 15], working memory [16, 17], attention [18], propagating waves [19, 20], non-rhythmic waking states [21, 22], or the emergence of up and down states [23]. It remains an open question how to compute realistically EEGs from such widely used network models of simple point neurons.

A major problem in achieving the above goal is that in such LIF point neurons all transmembrane currents collapse into a single point in space and the resulting extracellular potential is, therefore, zero [6]. Previous studies comparing the simulation output of networks of simple model neurons without a spatial structure with measures of graded extracellular potentials such as EEGs or LFPs have used ad-hoc approaches to estimate the EEG from variables available from simulation of the network, including the average membrane potentials [23–28], the average firing rate [29–31], the sum of all synaptic currents [13, 32, 33], or the sum of absolute values of synaptic currents [14, 34]. However, the limitations and caveats of using such ad-hoc simplifications to compute the EEG have been rarely considered and tested. As a result, it is still unclear how best to compute EEGs directly from output from point-like neuron network models [35, 36].

In order to generate extracellular potentials, spatially extended neuron models, i.e., multicompartment neuron models, are required [37, 38]. Previous studies have numerically computed the compound extracellular potential as the linear superposition of all single-cell distance-weighted transmembrane currents within a network of multicompartment neurons [39–41]. This approach is however computationally cumbersome, and it does not allow an easily tractable and exhaustive analysis of the dynamics of such networks. One alternative could be using a hybrid scheme [30, 35, 42, 43] that projects the spike times generated by the LIF point-neuron network onto morphologically detailed 3D neuron models and then computing the electric field that the currents flowing through these 3D networks generate. This scheme provides a simplification by separating the study of the network dynamics (described by the point-neuron network model) from that of field generation (described by the multicompartment neuron model), but still requires running cumbersome multicompartment model simulations for each simulation of the LIF network.

In this article, we implemented a much simpler and lighter method to predict the EEG based simply on the variables available directly from simulation of a point-neuron network model (e.g., membrane potentials, spike times or synaptic currents of the neuron models). We constructed several different candidate approximations (termed proxies) of the EEG that can be computed from networks of LIF point neurons. We then evaluated how well each proxy reconstructed a ground-truth EEG obtained when the synaptic input currents of the LIF network were injected into an analogous three-dimensional (3D) network model of multi-compartmental neurons with realistic cell morphologies. This approach was shown to perform remarkably well in predicting the LFP [42], based on a specific weighted sum of synaptic currents from the point-neuron network model, for a specific network state (i.e., asynchronous irregular) of the LIF network model. However, the previously obtained LFP proxy did not include a head model that approximates the different geometries and electrical conductivities of the head necessary for computing a realistic EEG signal recorded by scalp electrodes. We thus derived a new proxy for the EEG that was validated against detailed simulations of the multicompartment model, investigating different cell morphologies, variations of distribution of presynaptic inputs, and changes in position of the recording electrode. Unlike previous studies which focused on approximations valid in specific network states [42], we also validated our proxies across the repertoire of network states displayed by recurrent network models, namely the asynchronous irregular (AI), synchronous irregular (SI), and synchronous regular (SR) [12] states, with different patterns of global oscillations and individual cell activity. We found that a new class of simple EEG proxies, based on a weighted sum of synaptic currents, outperformed previous approaches, including those optimized for predicting LFPs [14, 42]. The new EEG proxies closely captured both the temporal and spectral features of the EEG. We also provided a non-linear refinement using a convolutional neural network to estimate the EEG from synaptic currents, which yielded moderate improvements over the linear proxy at the expense of increasing complexity of the EEG estimation model.

## Results

### Computing the ground-truth EEG and EEG proxies

We investigated how to compute a simple but accurate approximation of the EEG (“EEG proxy” hereafter) that would be generated by the activity of a LIF point-neuron network if its neurons had a realistic spatial structure. We therefore first simulated a well-established model of a recurrent network of LIF point neurons. We then fed the spiking activity generated by the LIF point-neuron network into a realistic three-dimensional multicompartmental network model of a cortical layer and computed the EEG generated by this activity. We finally studied how to approximate this EEG simply by using the variables directly available from the simulation of the point-neuron network model.

The LIF point-neuron network was constructed using a well-established two-population (one excitatory and one inhibitory) model of a recurrent cortical circuit [12], illustrated in Fig 1 A. The network receives thalamic synaptic input that carry the sensory information and stimulus-unrelated inputs representing slow ongoing fluctuations of cortical activity. This network can generate a repertoire of different network states that map well into empirical observations of cortical dynamics [12, 44]. Fig 1 B shows, as an example, the asynchronous irregular spiking activity generated by a subset of the excitatory and inhibitory populations in response to a low firing rate of the thalamic input. We have shown in previous work that this model captured well (even more than 90% of the variance of empirical data) the dynamics of primary visual cortex under naturalistic stimulation [14, 34, 45].

**Fig 1.**
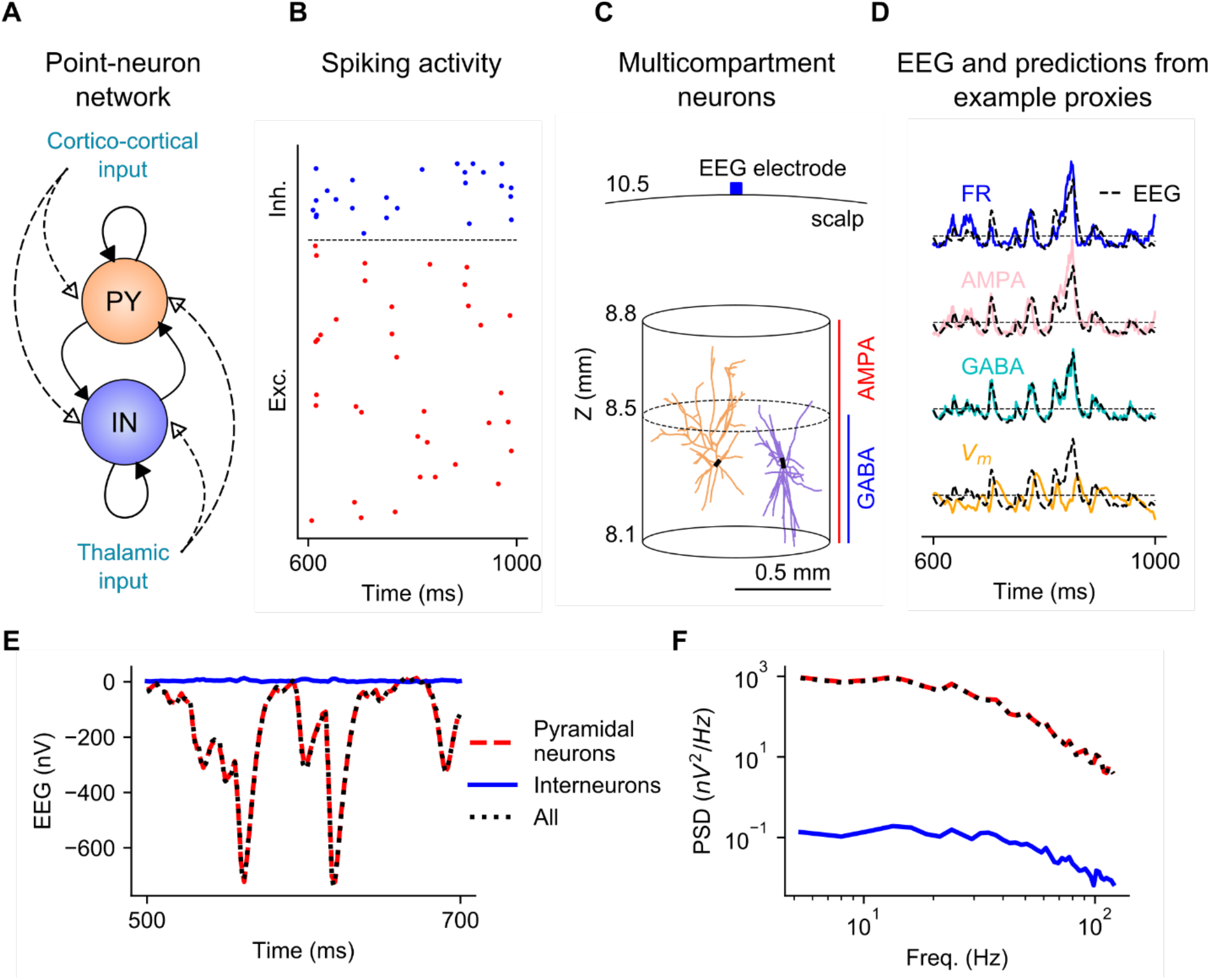
Overview of the network models and computation of proxies and EEG. (A) Sketch of the point-neuron network with recurrent connections between two types of populations: excitatory cells (pyramidal cells, PY) and inhibitory cells (interneurons, IN). Each population receives two kinds of external inputs: global ongoing cortico-cortical activity and thalamic stimulation. (B) Raster plot of spiking activity from a subset of cells in each population. (C) Sketch of the multicompartment neuron models used for generation of the EEG. Two representative model neurons are depicted, a pyramidal cell on the left and an interneuron on the right, positioned within a cylinder of *r* = 0.5 mm. While AMPA synapses are homogenously distributed over all compartments of both types of cells, GABA synapses on pyramidal cells are located only below *Z* = 8.5 mm. The EEG recording electrode is situated on the surface of the scalp layer. (D) Comparison between example proxies calculated from the point-neuron network and the ground-truth EEG computed from the multicompartment neuron model network. (E) EEG generated in the multicompartment neuron network by all neurons (dotted black), only pyramidal neurons (dashed red) or only interneurons (solid blue). (F) Corresponding power spectra for the three sets depicted in (E).

We then computed a “ground-truth” EEG (referred to simply as “EEG” in the paper), following the hybrid modelling scheme [30, 35, 42, 43], and used this ground-truth EEG to compare the performance of the different proxies. To do so, we created a network of unconnected multicompartment neuron models with realistic morphologies and homogeneous distribution within the circular section of a cylinder of radius *r* = 0.5 mm (Fig 1 C), which roughly approximates the spatial extension of a layer in a cortical column. We focused on computing the EEG generated by neurons with somas positioned in layer 2/3, so that somas of the multicompartment neurons are aligned in the Z-axis (150 μm below the reference point Z = 8.5 mm). We chose to position somas in layer 2/3 based on previous computational work suggesting that this layer gives a large contribution to extracellular potentials [30, 35]. The reference point Z = 8.5 mm was chosen to approximate the radial distance between the center of a spherical rodent head model and the brain tissue [46]. In this specific set of simulations performed for optimizing the proxies, we used the reconstructed morphology of a broad-tuft layer-2/3 pyramidal cell from rat somatosensory cortex available in the Neocortical Microcircuitry (NMC) portal [47, 48], referenced as *dend-C250500A-P3_axon-C260897C-P2-Clone_9* (see “Methods”). We chose this pyramidal-cell morphology because its open-field geometry is expected to generate large extracellular potentials. Inhibitory cells of the model were implemented using the morphology of L2/3 large basket cell interneurons (the most numerous class in L2/3 [47].

AMPA synapses were homogenously positioned along the Z-axis in both cell types, representing uniformly distributed excitatory input. In our default setting, we assumed that all inhibitory synapses are made by large basket cell interneurons of the model, which based on their morphology would be principally located below the reference point Z = 8.5 mm. Thus, all dendrites of inhibitory cells receive GABA synapses while only those dendrites of excitatory cells below Z = 8.5 mm receive GABA synapses, representing perisomatic inhibition.

EEGs were then generated from transmembrane currents of multicompartment neurons in combination with a forward-modelling scheme based on volume conduction theory [6]. To approximate the different geometries and electrical conductivities of the head, we computed the EEG using the four-layered spherical head model described in [35, 49]. In this model, the different layers represent the brain tissue, cerebrospinal fluid (CSF), skull, and scalp, with radii 9, 9.5, 10 and 10.5 mm respectively, which approximate the dimensions of a rodent head model [46]. The values of the chosen conductivities are the default values of 0.3, 1.5, 0.015 and 0.3 S/m. The simulated EEG electrode was placed on the scalp surface, at the top of the head model (Fig 1 C).

The time series of spikes of individual point neurons were finally mapped to synapse activation times on corresponding postsynaptic multicompartment neurons. Each multicompartment neuron was randomly assigned to a unique neuron in the point-neuron network and receives the same input spikes of the equivalent point neuron. Since the multicompartment neurons were not connected to each other, they were not involved in the network dynamics and their only role was to transform the spiking activity of the point-neuron network into a realistic estimate of the EEG. The EEG computed from the multicompartment neuron model network was then used as benchmark ground-truth data against which we compared different candidate proxies (Fig 1 D).

### Dynamic states of network activity of the point-neuron network model

The LIF point-neuron network model chosen to generate network dynamics is known to generate a number of qualitatively different activity states [12, 44] with patterns of variability of spike activity and network oscillations observed in cortical data. Since one of our goals is to determine EEG proxies which work well under a wide range of different network dynamics, we computed the different network states that the LIF point-neuron network can generate and which are recapitulated here.

The states generated by the LIF neuron network can be mapped by systematically varying across simulations the thalamic input (*v*_*0*_) and the relative strength of inhibitory synapses (*g*). We then use three different measures to describe the network dynamics: synchrony, irregularity, and mean firing rate [12, 44].

In Fig 2 A, we plot these three descriptors as a function of *g* and *v*_*0*_. We individuated 3 different regions of the parameter space, each corresponding to a qualitatively different network state, according to the criteria employed by Kumar and collaborators [44]. The asynchronous irregular (AI) state is characterized by a low value of network synchrony (< 0.01), an irregularity level close to the value of a Poisson generator (> 0.8) and a very low firing rate, below 2 spikes/s. The synchronous irregular (SI) state has a level of network synchrony higher than that of the AI state (between 0.01 and 0.1), but with highly irregular firing of individual neurons (irregularity above 0.8). In the SI, neurons spike at low rate (< 5 spikes/s). For the synchronous regular (SR) state, the network exhibits high synchronous activity (> 0.1), a more regular single-cell spiking (irregularity below 0.8) and high spiking rate (> 60 spikes/s). Spike raster plots of excitatory and inhibitory cell populations of representative samples selected for each network state are shown in Fig 2 B.

**Fig 2.**
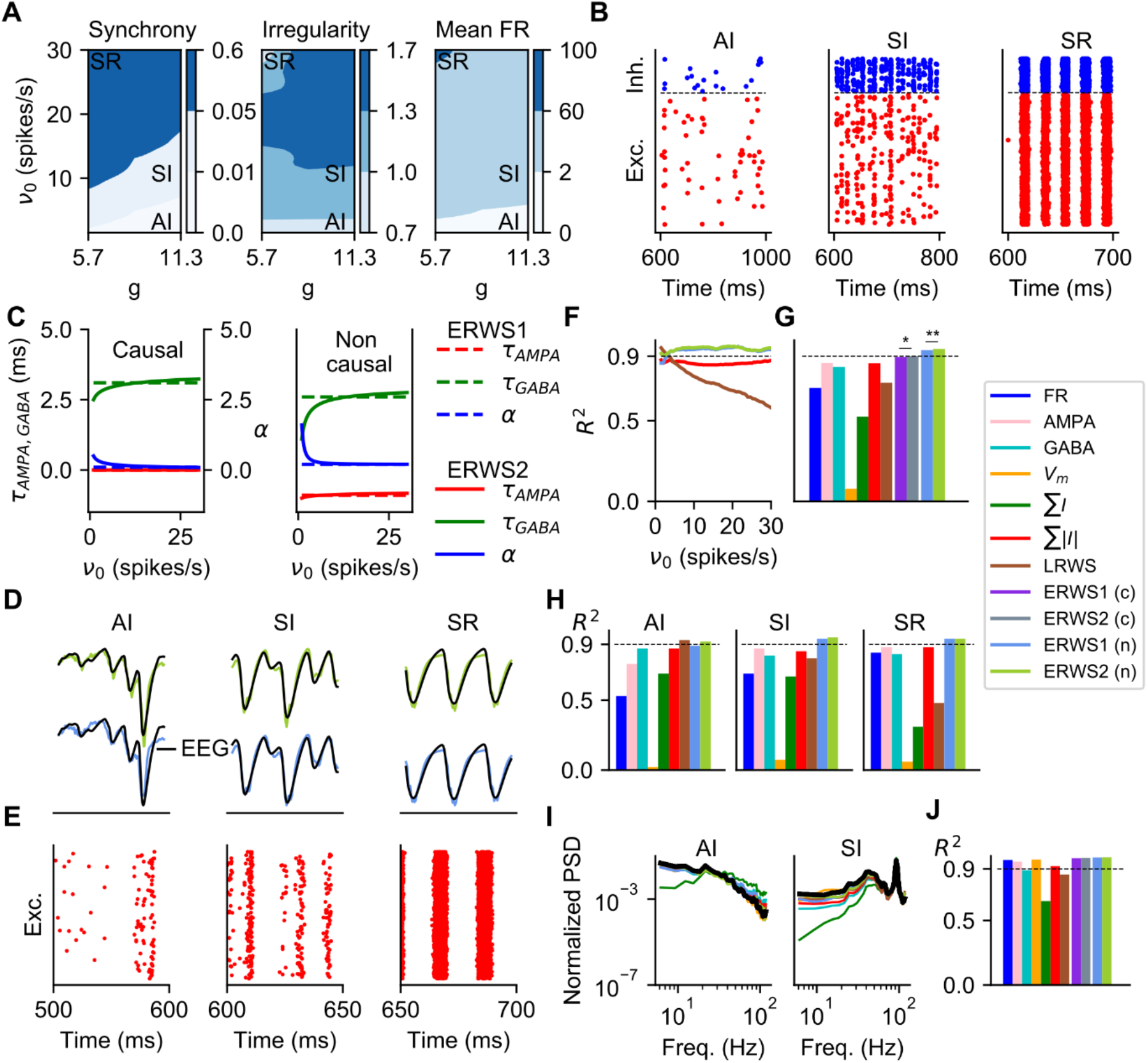
Optimization and validation of proxies for different sets of network parameters (*v*_*0*_, *g*). (A) Dynamic states of network activity defined by the control parameters *g* and *v*_*0*_. The labels AI (asynchronous irregular), SI (synchronous irregular) and SR (synchronous regular) indicate the combinations of parameters that have been selected as representative samples of each network state. The synchrony and irregularity are unitless, the mean firing rate (FR) is measured in spikes/s. (B) Spiking activity from a subset of cells of the excitatory and inhibitory populations for the same samples shown in (A). (C) Optimized parameters of *ERWS1* and *ERWS2* (Eqs. 7–9) as a function of the thalamic firing rate *v*_*0*_. We considered two alternative scenarios. In the causal version of the proxy, the output depends only on present and past inputs so that the time constants (*τ*_*AMPA*_ and *τ*_*GABA*_) are constrained to be positive. In contrast, non-causal proxies can be indifferently assigned positive and negative time constants. (D) Outputs of non-causal *ERWS1* (bottom row) and non-causal *ERWS2* (top row) proxies for different network states compared to ground-truth EEGs. (E) Spiking activity for the same simulation cases of panel D. (F) Average performance, evaluated by using the coefficient of determination *R*^*2*^, of ∑|*I*|, *LRWS*, *ERWS1* (non-causal) and *ERWS2* (non-causal) calculated on the validation dataset as a function of *v*_*0*_ (same colors as shown in (G)). (G) Average R^2^ of every proxy across all network instantiations *i* of the validation dataset (*c* is causal, *n* is non-causal). The same colors shown in this legend are used throughout the article to identify the different proxies. Tests for statistical significance are computed only for the pair *ERWS1* (non-causal) and *ERWS2* (non-causal) and for the pair *ERWS1* (causal) and *ERWS2* (causal). (H) R^2^ across network states. (I) Power spectral density (PSD) of the proxies and the EEG (in black). (J) Average R^2^ applied to proxies’ PSDs instead of their temporal responses. R^2^ is computed in the 5-200 Hz frequency range.

### Optimization and validation of proxies across different network states

We investigated how best to compute the proxy that combines the variables available directly from the simulation of a LIF point-neuron network model for accurately predicting the EEG over a wide range of network activity states. We explored different proxies that have been commonly used in previous literature for estimating the extracellular signal from point-neuron networks: (i) the average firing rate (*FR*), (ii) the average membrane potential (*V*_*m*_), (iii) the average sum of AMPA currents (*AMPA*), (iv) the average sum of GABA currents (*GABA*), (v) the average sum of synaptic currents (∑*I*) and (vi) the average sum of their absolute values (∑|*I*|). Furthermore, we propose here a new class of current-based proxies, (vii) the EEG reference weighted sum 1 (*ERWS1*) and (viii) the EEG reference weighted sum 2 (*ERWS2*), which are optimized linear combinations of time-delayed measures of AMPA and GABA currents. Indeed, an optimized weighted sum of synaptic currents (defined here as *LRWS*) was previously shown to be a robust proxy for the LFP [42]. The difference between *ERWS1* and *ERSW2* is that parameters of *ERWS2* adapt theirs values as a function of the strength of the external thalamic input *u*_0_, whereas the parameters of *ERWS1* are not dependent on *u*_0_ (see “Methods”).

We only considered the transmembrane currents of pyramidal cells to generate the EEG (in the multicompartment neuron network) because the contribution of transmembrane currents of interneurons to the EEG was shown to be negligible (Fig 1 E and F), in line with findings of Refs. [35] for the EEG and [42] for the LFP. Interneurons, though, play an indirect role in generating the EEG, since GABAergic currents in pyramidal cells depend on interneuronal spikes. In a similar way, proxies of the LIF neuron network are computed only on excitatory neurons.

The firing rate of inhibitory neurons might be expected to contribute as well to the *FR* proxy and, as a consequence, to the EEG, as observed in [30]. To keep consistency with definition of the other proxies, we decided to compute the *FR* proxy based only on firing rates of excitatory cells. We checked that using a proxy computed on firing rates of both excitatory and inhibitory cells gave an EEG reconstruction accuracy considerably poorer than accuracy of the proxies based on synaptic currents (from proxy *iii* to proxy *viii* above).

The first 6 proxies taken from previous literature are parameter-free. The two new ones, *ERWS1* and *ERWS2* have 3 and 9 free parameters, respectively, which need to be optimized (Eqs. 7–9). Following previous work [42], these parameters define the factor *α* describing the relative ratio between the two currents and a specific delay for each type of current (*τ*_*AMPA*_, *τ*_*GABA*_). We computed the values of these parameters by a cross-validated optimization of the predicted EEG across the different network states seen for the LIF network model.

For optimization and validation of proxies we generated a large set of numerical simulations (522 simulations) by systematically varying the values of *g* and *v*_*0*_ over a wide state range. In each simulation instantiation, we set a given value *g* and *v*_*0*_ and used different random initial conditions (e.g., recurrent connections of the point-neuron network or soma positions of multicompartment neurons). The best-fit values of *ERWS1* and *ERWS2* were calculated by minimizing the sum of square errors between the ground-truth EEG and the proxy for all network instantiations of the optimization dataset (see “Methods”, Eq. 11).

Fig 2 C shows the best parameters (*α*, *τ*_*AMPA*_ and *τ*_*GABA*_) found by the optimization algorithm for the two alternative scenarios considered here: causal and non-causal proxies (see also Table 1). For causal proxies, the predicted EEG depended only on present and past values of AMPA and GABA currents. Thus, the time delay parameters *τ*_*AMPA*_ and *τ*_*GABA*_ (quantifying the delay by which the synaptic current contributes to the EEG) were constrained during optimization to be non-negative. For non-causal proxies, time delay parameters can take positive and negative values. Non-causal relationships between measured extracellular potentials and neural activity at multiple sites may emerge because of closed-loop recurrent interactions within the network [6]. The mathematical expressions of the optimized causal proxies are:

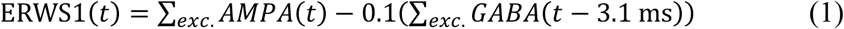

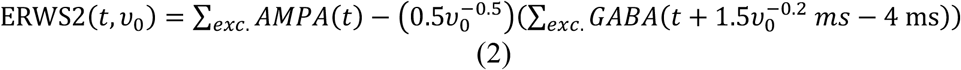

**Table 1.**
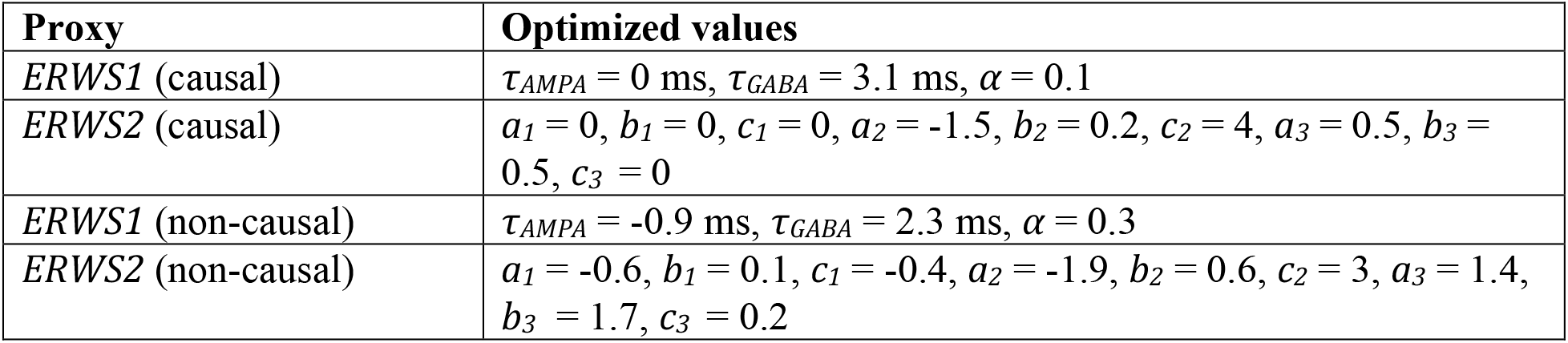
Parameters of *ERWS1* and *ERWS2*.

| Proxy | Optimized values |
| --- | --- |
| *ERWS1* (causal) | $\tau_{AMPA} = 0$ ms, $\tau_{GABA} = 3.1$ ms, $\alpha = 0.1$ |
| *ERWS2* (causal) | $a_1 = 0$, $b_1 = 0$, $c_1 = 0$, $a_2 = -1.5$, $b_2 = 0.2$, $c_2 = 4$, $a_3 = 0.5$, $b_3 = 0.5$, $c_3 = 0$ |
| *ERWS1* (non-causal) | $\tau_{AMPA} = -0.9$ ms, $\tau_{GABA} = 2.3$ ms, $\alpha = 0.3$ |
| *ERWS2* (non-causal) | $a_1 = -0.6$, $b_1 = 0.1$, $c_1 = -0.4$, $a_2 = -1.9$, $b_2 = 0.6$, $c_2 = 3$, $a_3 = 1.4$, $b_3 = 1.7$, $c_3 = 0.2$ |

Expressions of the optimized non-causal proxies (*v*_*0*_ is unitless) are:

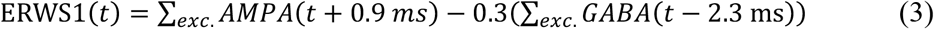

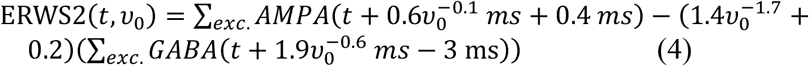

We first show the best fits obtained from optimization of the two *ERSW* proxies (Fig 2 C). For both *ERWS1* and *ERWS2*, in the non-causal versions, the time delay parameters were small (few milliseconds) but had opposite signs, *τ*_*GABA*_ was positive while *τ*_*AMPA*_ was negative. In the causal version of both proxies, we observed a similar trend but *τ*_*AMPA*_ was constrained to 0 by the optimization. Thus, the best EEG proxies depend on past values of GABA synaptic currents and on current and future values of AMPA synaptic currents. These values are different from the optimal delays (*τ*_*GABA*_ = 0 ms and *τ*_*AMPA*_ = 6 ms) found for the LFP in [42]. One reason for the observed difference between the previous LFP proxy and our new EEG proxies may relate to differences in spatial integration properties of the EEG signal and the LFP signal. Another probable cause of this difference is that in [42] the LFP proxy was optimized over a much smaller range of network states and external input rates (*v*_*0*_ < 6 spikes/s). Indeed, our results for *ERWS2* show that optimal values of *τ*_*GABA*_ exhibit strong adaptation towards *τ*_*GABA*_ = 0 ms within the low regime of the external rate *v*_*0*_. The parameter *α*, which expresses the ratio of the contribution to the EEG of GABA relative to AMPA synaptic currents, also exhibits a strong adaptation. The dependence of *α* on the value of input rate *v*_*0*_ in Fig 2 C is particularly relevant because it reflects a larger weight of GABA currents for low values of *v*_*0*_ and the opposite effect, stronger weight of AMPA currents, as the external rate increases.

To quantitatively evaluate the performance of all proxies, we computed for each proxy the coefficient of determination *R*^*2*^, which represents the fraction of the EEG variance explained. The average *R*^*2*^ calculated on the validation dataset (Fig 2 G) shows a clear superiority of the new class of proxies. Both the causal and non-causal versions of *ERWS1* and *ERWS2* outperform all the other proxies, and the non-causal versions reach the best overall performance (*ERWS1: R^2^* = 0.94 and *ERWS2: R^2^* = 0.95). In agreement with previous results for the LFP [42], the three proxies that give the worst fits were *FR*, ∑*I* and *V*_*m*_.

To understand if the performance of proxies depended on the specific state of network activity, we first examined the performance of the most interesting proxies (∑|*I*|, *LRWS*, *ERWS1* (non-causal) and *ERWS2* (non-causal)) separately for different values of the input rate *v*_*0*_. We found that while *LRWS* performs well for low input rates (the range of external rates for which it was optimized [42]), its performance rapidly dropped with *v*_*0*_ (Fig 2 F). The other three proxies maintained a high *R*^*2*^ for the whole spectrum of firing rates studied here, with *ERWS1* and *ERWS2* performing notably better than ∑|*I*|. Note also that *ERWS2* is the only proxy that yields a value of *R*^*2*^ above 0.9 for all firing rates. We then computed the performance of these proxies separately for different types of network states. We found that the new proxies developed here, *ERWS1* and *ERWS2*, produced accurate fits of the EEG for all network states (Fig 2 H), while accuracy of EEG approximations made by the other proxies was less uniform across network states.

The above analyses quantified how well the proxies approximated the actual values of the EEG in the time domain for each data point. We next examined how well the proxies approximated the overall power spectrum of the EEG rather than all variations of the EEG time series. In Fig 2 I we show power spectral density (PSD) functions of all the proxies for the AI and SI states, compared to spectral responses of the EEG. In the whole frequency range considered (5 – 200 Hz), all proxies provide a prominent good fit of the EEG power spectrum, except ∑*I*, which attenuates low frequencies and amplifies high frequencies. In Fig 2 J we report the average R^2^ computed for the PSDs across all data points of the validation dataset, confirming that all proxies gave an accurate approximation of the EEG power spectrum (except ∑*I*).

In sum, while almost all proxies are good enough to capture the general properties of the EEG power spectrum, *ERWS1* and *ERWS2* capture best the details of time variations of the EEG.

### Time-shifted variants of proxies

The *ERSW* proxies were optimized for EEG prediction choosing optimal values for the time shifts between neural activity and the EEG. It is thus possible that the superior performance of the *ERWS* proxies over all others may have been due to the fact that the other proxies were not optimally time shifted. To investigate this hypothesis, we generated optimized time-shifted versions of all the other proxies by computing cross-correlation between the ground-truth EEG and all other proxies and choosing the optimum time shift of each proxy as the lag of the cross-correlation peak. We then compared the performance of the time-shifted versions of proxies in predicting the EEG with the performance of the *ERWS* proxies.

In this analysis, we recomputed the optimum time shift of every proxy separately for each network state, whereas the parameters of the *ERWS* proxies were jointly optimized (see previous section) over the entire simulated EEG dataset spanning all possible network states. Thus, this comparison was clearly favorable to the other proxies. Nevertheless, we still found that the *ERWS* proxies outperformed all previous proxies for the majority of network states. Only in the AI state, we observed that the *LRWS* proxy slightly outperformed *ERWS1* and *ERWS2*. The *ERWS2* proxy was the only one providing remarkably good performance across all states (*R*^*2*^ > 0.9 over all states).

Further results came out of this analysis. Two proxies clearly improved the quality of their fits after time shifting, *FR* and *V*_*m*_, but presented opposed time shifts: while *FR* was delayed, *V*_*m*_ was moved forward in time. A spike is a local and instantaneous event in time and, as a result, a firing-rate proxy is expected to exhibit faster temporal changes than the EEG signal. By contrast, integration of the postsynaptic soma membrane potentials following presynaptic spiking is a slower process that might lead to a signal more low-pass filtered than the EEG.

When comparing AMPA and GABA proxies, we observed that, in the AI state (Fig 3 A), temporal dynamics of the EEG signal were better approximated by the GABA proxy, whereas AMPA currents showed a faster response. Indeed, the performance of the *AMPA* proxy was improved after applying the corresponding time shift. As the firing rate of the external input increased and switched the network state from AI to SI (Fig 3 B), the temporal evolution of the EEG began to diverge from GABA currents and, instead, AMPA currents were seen to better approximate the EEG.

**Fig 3.**
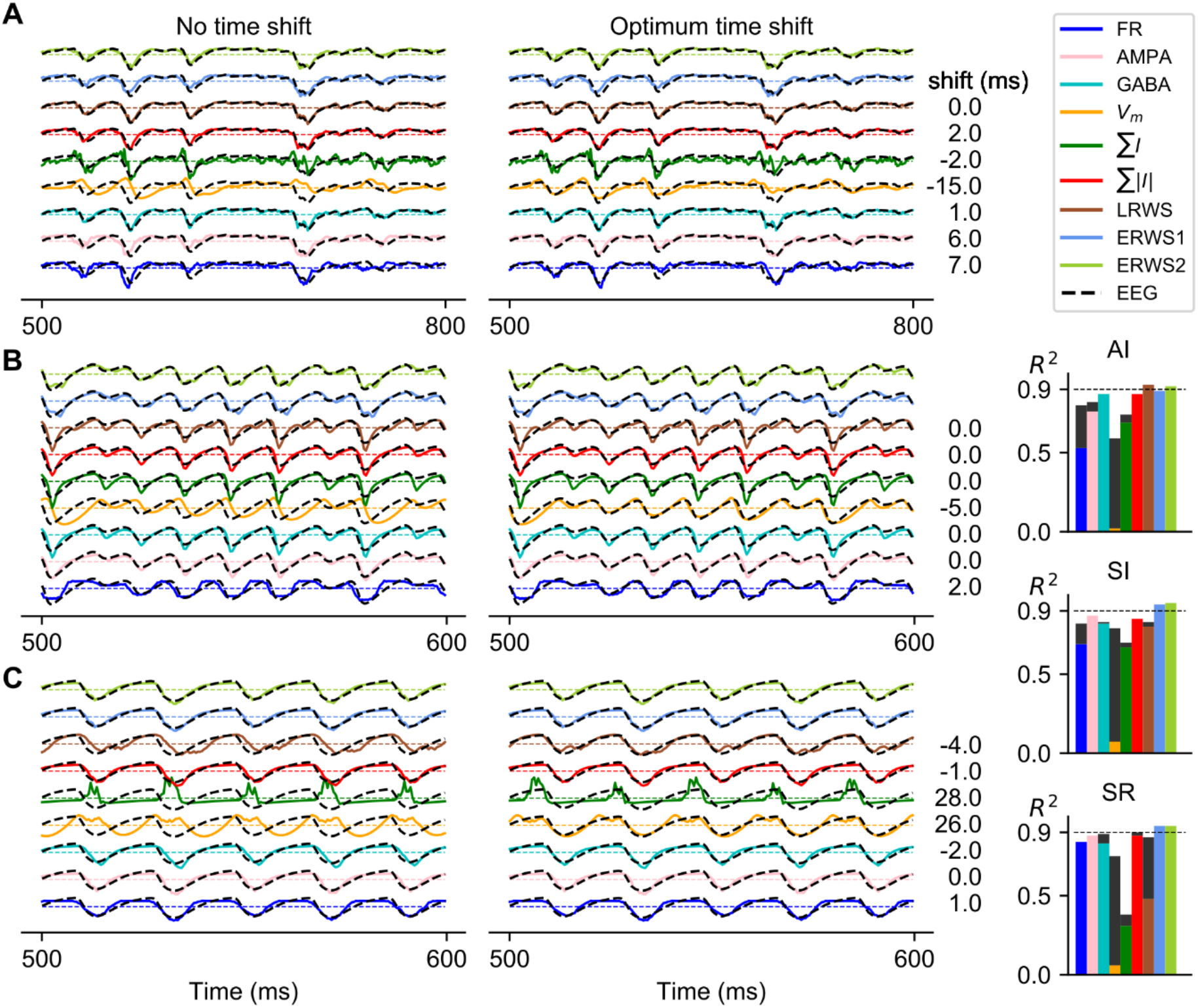
Optimum time shift of proxies that maximizes cross-correlation with the EEG. Comparison of the outputs of proxies and the ground-truth EEG before (left) and after (right) applying the optimum time shift, with the optimum time shift for each proxy and network state indicated on the right. Note that some proxies have positive time shifts for all network states (e.g., *FR*), while others (e.g., *GABA*) change the sign of the time shift when passing from the AI to the SR state. The network states shown are the following: AI in panel A, SI in panel B and SR in panel C. On the right: *R*^*2*^ before (color bars) and after (black bars) applying the optimum time shift. *ERWS1* and *ERWS2* are not time shifted.

### The performance of EEG proxies depends on the neuron morphology and distribution of synapses

Modelling studies have demonstrated that extracellular potentials generated by synaptic input currents vary with the neurons’ dendritic morphology and the positions of individual synaptic inputs [6, 50]. For example, morphological types that display a so-called *open-field* structure, such as pyramidal cells, have spatially separated current sources and current sinks that generate a sizable current dipole. Synaptic inputs onto neurons that have a *closed-field* configuration, such as interneurons, largely cancel out when they are superimposed so that the net contribution to the current dipole is weak [35]. The hybrid modelling scheme [30, 35, 42, 43] gives us the opportunity to study, independently from the spiking dynamics of the point-neuron network, how different parameters of the multicompartment neuron network (e.g., distribution of synapses or dendritic morphology) affect the EEG signal and, as a consequence, modify the prediction capabilities of the proxies.

Above results (Figs. 2 and 3) were computed using a specific multicompartmental model type of L2/3 pyramidal cell from rat somatosensory cortex (taken from the NMC database [47, 48]) and referred as “NMC L2/3 PY, clone 9” (Table 5, Figure 4A). Here, we studied whether the proxies derived for this morphology provided good approximations to the EEG generated by different cell morphologies. We thus quantified how well our proxies approximate the EEG generated by a different pyramidal-cell morphology taken also from rat somatosensory cortex (“NMC L2/3 PY, clone 0”) and by a third morphology (“ABA L2/3 PY”), which is a L2/3 pyramidal cell from the mouse primary visual area [51]. It is important to note that the parameter values of proxies optimized for the morphology “NMC L2/3 PY, clone 9” were applied unchanged to the other morphologies across network states.

**Fig 4.**
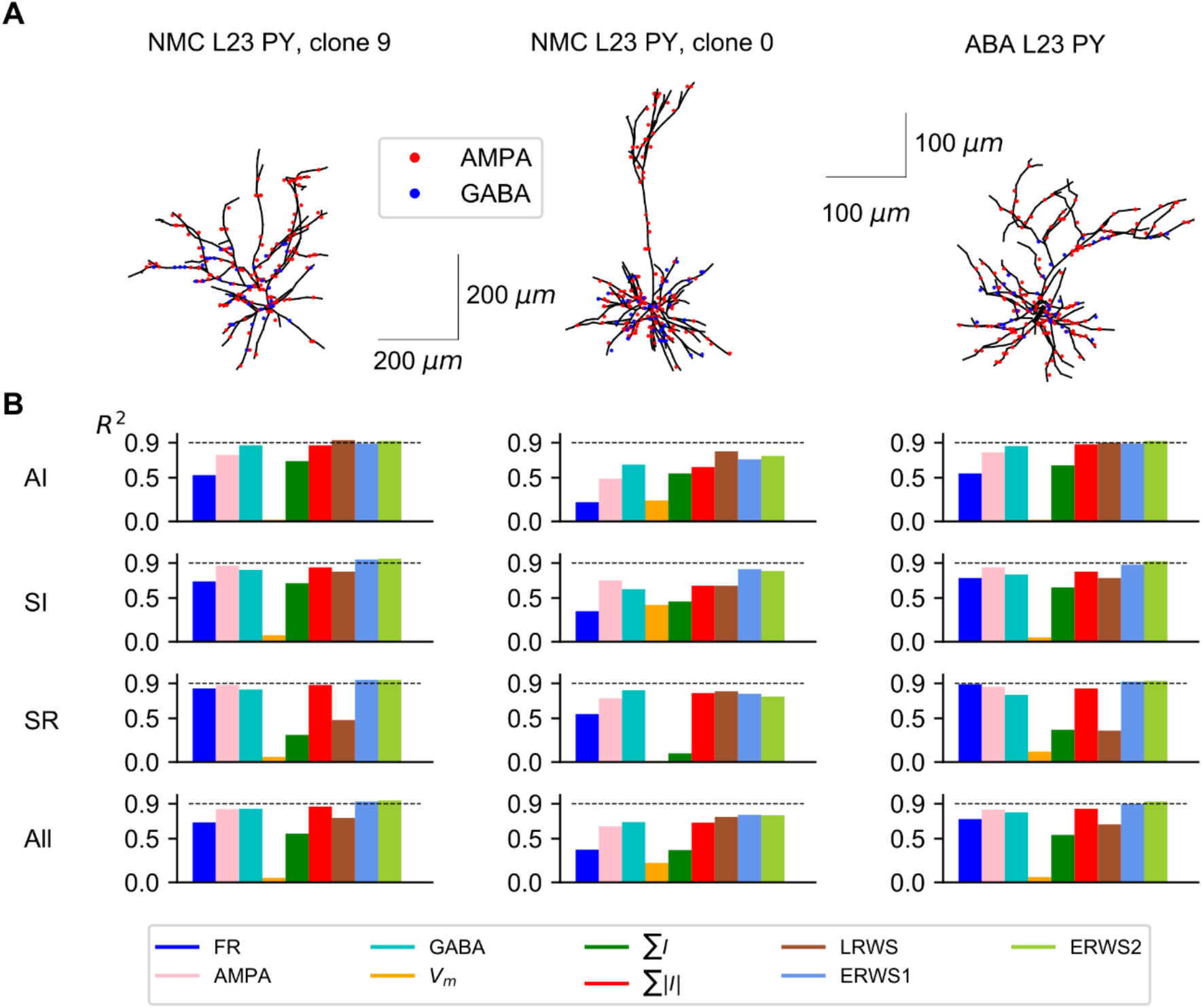
Performance of proxies for different morphologies. (A) Neuron reconstructions of L2/3 pyramidal cells acquired from the Neocortical Microcircuitry (NMC) portal [47, 48] and the Allen Brain Atlas (ABA) [51] (Table 5). For visualization purposes, in the synaptic distribution of each morphology, only a subset of AMPA and GABA synapses are shown, drawn randomly from all presynaptic connections. (B) *R*^*2*^ computed for each morphology (columns) and network state (rows). The label “All” indicates the average *R*^*2*^ across the three network states.

We found that *ERWS2* was the proxy with the highest prediction accuracy (Fig 4). It approximated extremely well the EEG across all three types of morphology and across all network states. The performance of both *ERWS* proxies in predicting the EEG generated by the mouse pyramidal neuron morphology (“ABA L2/3 PY”, Fig 4, right column) was as good as the performance for the “NMC L2/3 PY, clone 9” morphology (probably because they have similar broad-tuft dendritic morphology, although different size). This suggests that the model generalizes reasonably well across species (at least for EEG generated by broad-tuft dendritic morphologies). *ERWS* proxies also performed well, though less compared to the morphology they were optimized for, on the EEGs generated by the other rat somatosensory cortex morphology (“NMC L2/3 PY, clone 0”, Fig 4, middle column). The small decrease in performance was probably due to the fact that, unlike the broad dendritic tuft morphology used to optimized the proxy, this morphology incorporates long apical dendrites that separates AMPA synapses located in the tuft from GABA synapses more than 200 μm. Analogously, the similarity in performance of *LRWS* for the “NMC L2/3 PY, clone 0” morphology could be understood in terms of similarity between the pyramidal-cell morphology used to develop the *LRWS* proxy [42] and this morphology. The *LRWS* proxy [42] performed well across all morphologies in the AI state but its performance decreased across other states and morphologies. Other proxies performed poorly across different morphologies and/or states.

We next investigated how different spatial distributions of synapses on excitatory cells affect the performance of proxies (Fig 5). More specifically, GABA synapses were distributed on excitatory cells following two alternative approaches: located only on the lower part of the cell, primarily on the soma and basal dendrites (“Asymmetric”) or homogeneously distributed across all dendrites (“Homogeneous”). Note that the “Asymmetric” case (Fig 5, left column) corresponds to default configuration shown in Fig 4 A, left column (“NMC L2/3 PY, clone 9” morphology). The most significant change observed when distributing GABA synapses homogeneously on excitatory cells was an overall decrease of the performance of all proxies (but see ∑*I*), most prominently for the AI. These findings are in agreement with previous results obtained for the LFP proxy [42] in which an homogenous distribution of AMPA and GABA synapses on pyramidal cells resulted in the worst approximation of LFPs. In all scenarios, except for the AI state, *ERWS1* and *ERWS2* provided the best performance and their average *R*^*2*^ values across network states reflect their superiority in both the asymmetric and homogenous distributions.

**Fig 5.**
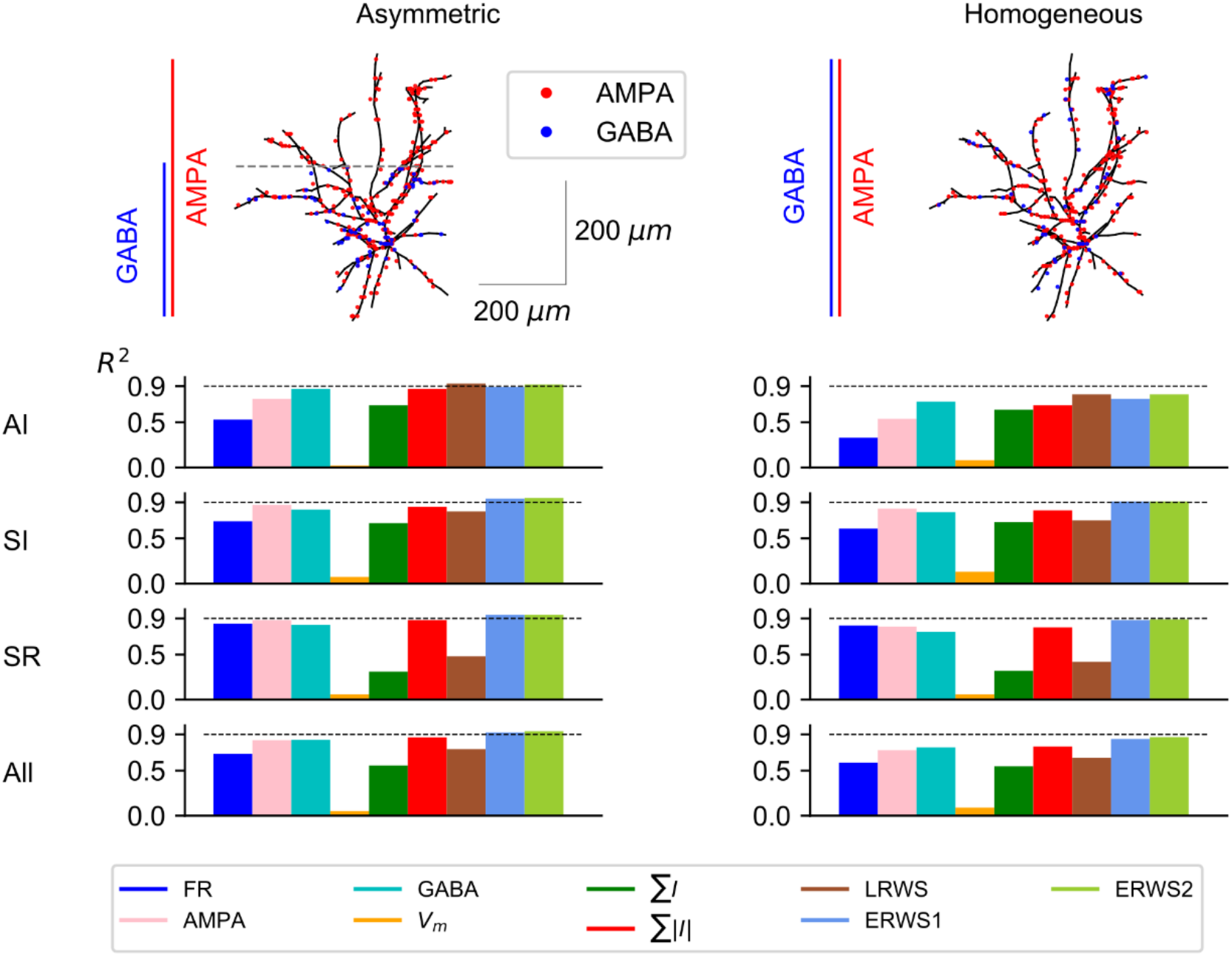
Influence of synaptic distributions on performance of proxies. Outline of the two different distributions of GABA synapses on excitatory cells: distributed only below the reference point *Z* = 8.5 mm (“Asymmetric”) or distributed homogenously across all dendrites (“Homogeneous”). Each row below the diagram of model cells shows the corresponding *R*^*2*^ for a different network state. The label “All” in the last row displays the average *R*^*2*^ across the three network states.

### Effects of the position of the electrode over the head model on the EEG and proxies

To investigate the relationship between the position of the electrode and its effects on the EEG and performance of proxies, we simulated the EEG at four different locations over the head (Fig 6 A). Simulation results are shown as a function of the angle between the electrode location and the Z-axis (Theta), computed for the three different network states: AI, SI and SR. We first explored how properties of the EEG signal changed with the location of the electrode. As expected, the EEG amplitude, defined as the standard deviation of the EEG signal over time, decreased steeply when the electrode is moved away from the top of the head (Fig 6 B). This decrease in EEG amplitude is consistent with previous simulation results of the 4-sphere head-model [35, 39], in which a moderate attenuation of the EEG scalp potentials was observed when increasing the lateral distance from the center position along the head surface. Although the EEG amplitude is larger in the SR state, the relative variations of amplitude as a function of Theta were similar across network states. In contrast, we found (Fig 6C) sizeable differences in the normalized time courses of the EEG at different network states: an increase of Theta involved a delay of the EEG signal that is larger for the AI and SI states, but much weaker for the SR state. These results could indicate that as the measurement point moves toward the zero-region of the current dipole, where the EEG power is much smaller, the signal-to-noise ratio is reduced and the influence of the high-frequency noise is more important. Since the signal power is significantly larger for the SR state, the effects of the high-frequency noise are less evident for the SR state.

**Fig 6.**
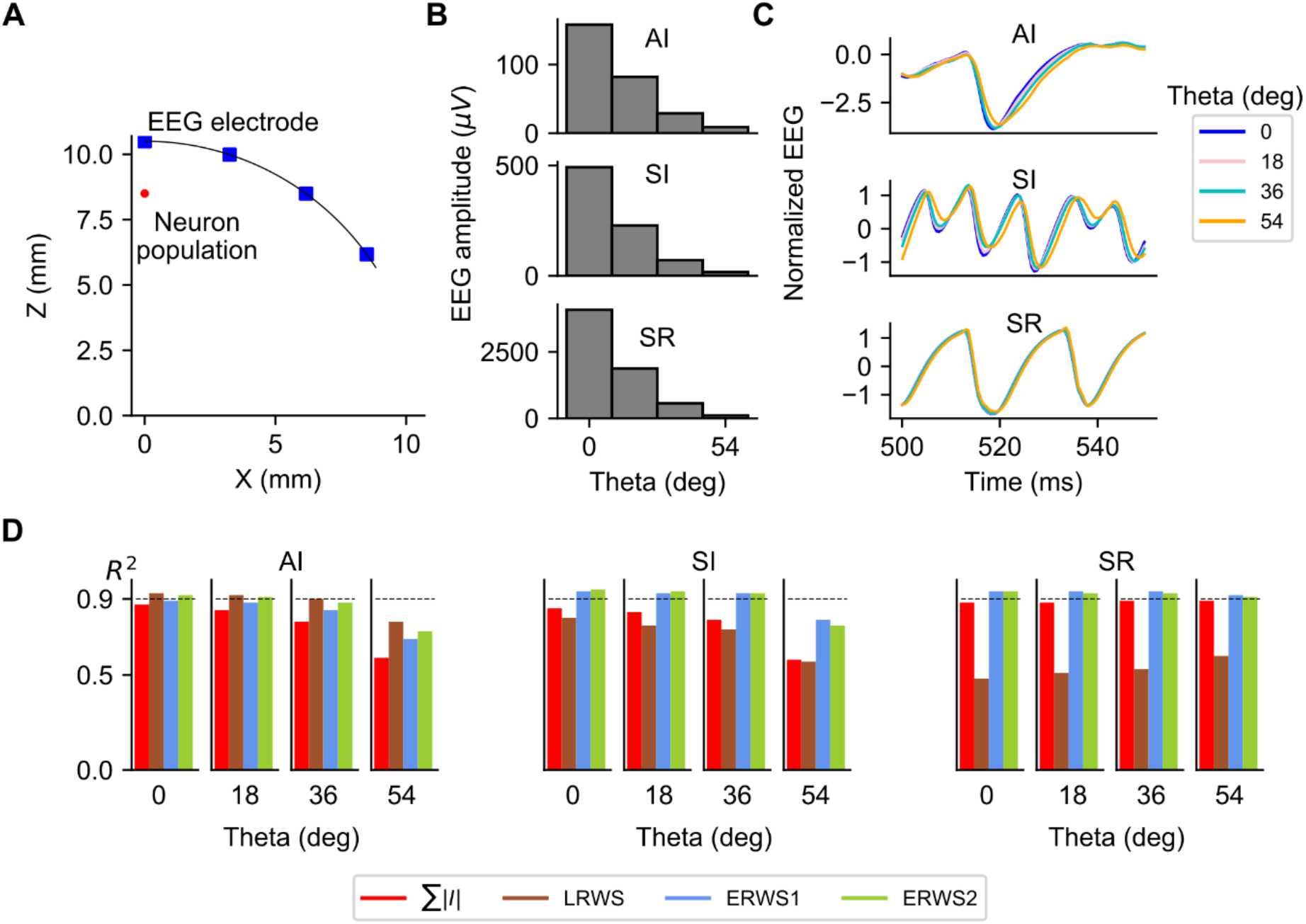
EEG and proxies as a function of the position of the electrode over the head model. (A) Illustration of the scalp layer in the four-sphere head model and locations where the EEG is computed. Location of the center of soma positions of the multicompartment neurons is marked as “Neuron population”. (B) EEG amplitude, (C) normalized EEG and (D) performance of ∑|*I*|, *LRWS*, *ERWS1* and *ERWS2* as a function of the angle between the electrode location and the Z-axis (Theta), computed for the three different network states: AI, SI and SR.

Variations of properties of the EEG signal when the electrode was shifted from the top of the head affected the performance of proxies. As depicted in Fig 6 D, the performance of ∑|*I*|, *LRWS*, *ERWS1* and *ERWS2* decreased when Theta was augmented in the AI and SI states. However, the performance of proxies is hardly modified by the position of the electrode in the SR state, or it even shows the opposite trend (an increase) in the case of the *LRWS* proxy. In any case, *ERWS1* and *ERWS2* give the best performance in most scenarios, particularly *ERWS2* whose *R*^*2*^ value is above 0.9, provided that Theta is smaller than 36 degrees.

### EEG estimation by CNN

The proxies considered above are all simple linear functions of the neural parameters of the LIF point-neuron network model. Linear proxies have the advantage of simplicity and interpretability. However, an alternative strategy for constructing an EEG proxy is training a convolutional neural network (CNN) to learn complex and possibly non-linear relationships between parameters of the LIF point-neuron network model, such as AMPA and GABA currents, and the EEG. This could potentially improve the estimation of linear proxies, at a possible expense of increasing computational complexity and hindering interpretation. Instead of using a deep neural network with many hidden layers that could largely increase complexity and prevent us from making any type of analogy with results of linear proxies, we opted for a simpler, shallow CNN architecture, with just one convolutional layer (Fig 7 A). This CNN architecture was found to be sufficiently robust achieving a *R*^*2*^ value of 0.99 on the test dataset (see Table 2). The network consists of one 1D convolutional layer (‘Conv1D’) with 50 filters and a kernel of size 20, followed by a max pooling layer (‘MaxPooling1D’) of pool size 2, a flatten layer and two fully connected layers of 200 units each one (marked as ‘Dense’ and ‘Output’ respectively). The input of the CNN is constructed by stacking data chunks of 100 ms (0.5 ms time resolution) extracted from the time series of AMPA and GABA currents, giving a 2 x 200 input layer.

**Fig 7.**
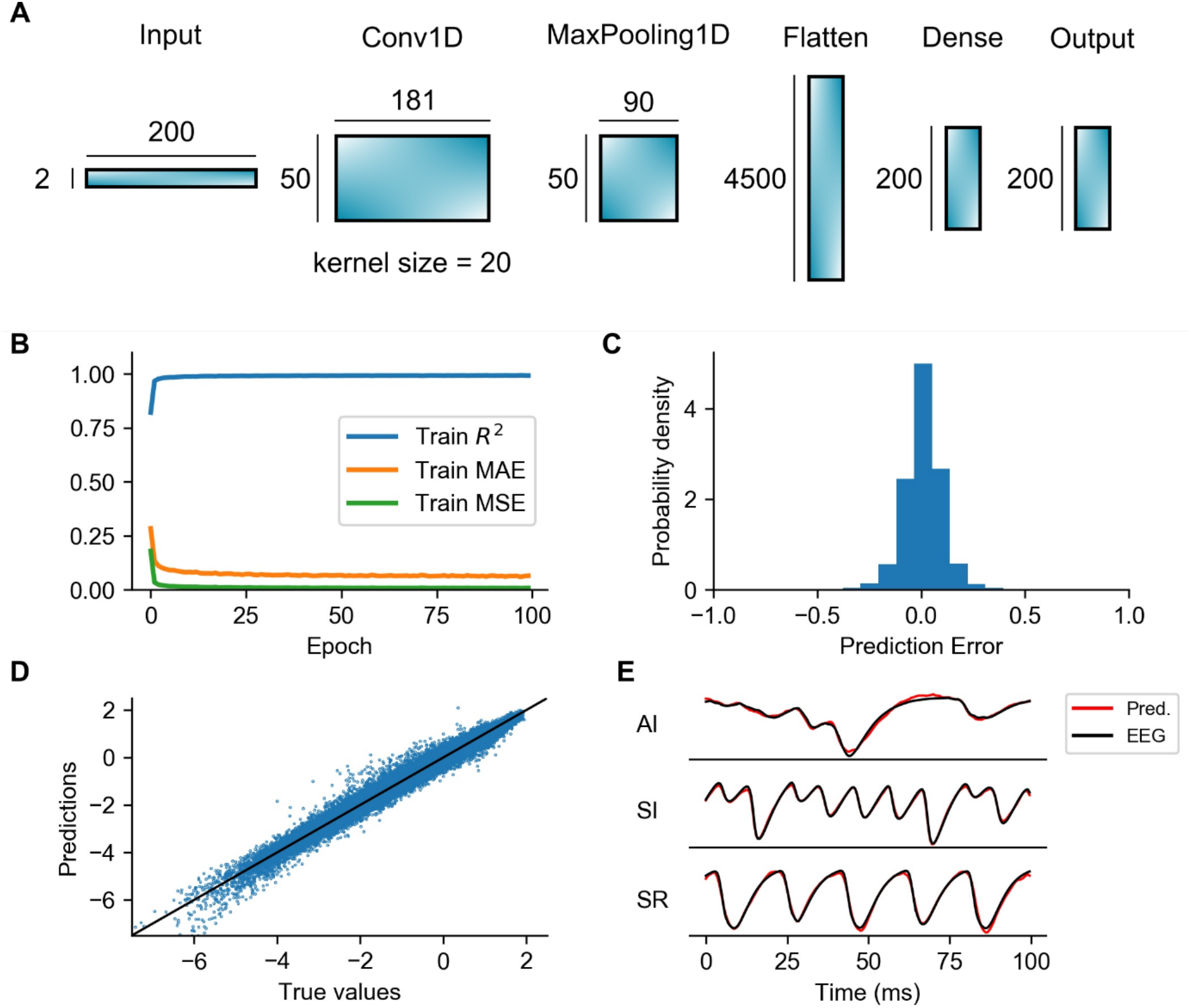
Overview of the convolutional neural network, train errors and accuracy of EEG predictions. (A) Illustration of the different types of layers included in the processing pipeline of the CNN architecture as well as the output shapes of each layer. Note that the 1D convolutional layer (‘Conv1D’) uses 50 filters and a 1D convolutional window (kernel) of size = 20. The total number of parameters of the entire CNN is 942450. (B) Training metrics collected during training: *R*^*2*^, Mean Absolute Error (MAE) and Mean Squared Error (MSE). (C) Probability density function of the prediction error calculated on the test dataset. (D) Predictions vs true values. Each dot of the scatter plot corresponds to amplitude values of the predicted and real EEG signals at a specific time step of the simulation. The continuous line represents a perfect EEG estimator. (E) Examples of predictions of the CNN compared to the ground-truth EEGs for different network states.

**Table 2.** Performance (computed as *R*^*2*^) of the CNN in comparison with ∑|*I*|, *LRWS, ERWS1* and *ERWS2* proxies. The performance values shown for the test dataset (A) are averaged over all samples of the test dataset, while performance values in panels B, C and D are averaged over the samples of the different network states, i.e., AI, SI and SR.

| A: Performance on the test dataset |  |  |  |  |  |
| --- | --- | --- | --- | --- | --- |
| $\Sigma\vert I\vert $ | *LRWS* | *ERWS1* | *ERWS2* | CNN | |
| 0.86 | 0.74 | 0.94 | 0.95 | 0.99 |  |
| B: Morphologies |  |  |  |  |  |
| Cell model | $\Sigma\vert I\vert $ | *LRWS* | *ERWS1* | *ERWS2* | CNN |
| NMC L2/3 PY, c. 9 | 0.87 | 0.74 | 0.92 | 0.94 | 0.97 |
| NMC L2/3 PY, c. 0 | 0.70 | 0.76 | 0.77 | 0.77 | 0.87 |
| ABA L2/3 PY | 0.85 | 0.67 | 0.90 | 0.92 | 0.94 |
| **C: Distribution of synapses** |  |  |  |  |  |
| **Distribution type** | **$\Sigma\vert I\vert $** | ***LRWS*** | ***ERWS1*** | ***ERWS2*** | **CNN** |
| Asymmetric | 0.87 | 0.74 | 0.92 | 0.94 | 0.97 |
| Homogeneous | 0.77 | 0.65 | 0.83 | 0.87 | 0.89 |
| **D: Position of the EEG electrode** |  |  |  |  |  |
| **Theta (rad)** | **$\Sigma\vert I\vert $** | ***LRWS*** | ***ERWS1*** | ***ERWS2*** | **CNN** |
| 0 | 0.87 | 0.74 | 0.92 | 0.94 | 0.97 |
| 0.31 | 0.86 | 0.74 | 0.91 | 0.93 | 0.97 |
| 0.63 | 0.82 | 0.72 | 0.90 | 0.91 | 0.96 |
| 0.94 | 0.69 | 0.68 | 0.80 | 0.81 | 0.87 |

The network was trained and tested on the same independent datasets (one for the training of the proxies, the other for the validation/testing of their accuracy) generated for optimization of parameters of the *ERWS1* and *ERWS2* proxies, using a first-order gradient descent method (Adam optimizer [52]) over 100 epochs (see Methods). In Fig 7 B, we observe a quick convergence of the three metrics used to monitor training (*R*^*2*^, MAE and MSE) towards optimal values (*R*^*2*^ ≈ 1, MAE < 0.1 and MSE < 0.01). Accuracy of predictions of the trained network, calculated on the test dataset, are shown in Fig 7 C-E. The probability distribution of the prediction error is depicted in panel C. Here we define the prediction error as the difference between amplitude values of the predicted and true EEG signals at a specific time step of the simulation. As observed, the prediction error distribution approximates a normal distribution with zero mean and standard deviation ≈ 0.1. The scatter plot of true versus predicted values (panel D) generally reflects a very accurate estimation of the EEG values with the swarm of points showing a clear trend that closely follows the line of a perfect EEG estimator. In panel E, we illustrate some examples of predictions of the EEG signal compared to the ground-truth EEG for different network states. Interestingly, the best match between predicted and true EEG traces is seen for the SI state, although the other two states, AI and SR, produce also fairly good estimations.

The performance of the CNN was evaluated, like for the other proxies, as the average value of *R*^*2*^ computed over all samples of the test dataset. As shown in Table 2 A, the CNN clearly outperformed all other proxies on the test dataset and reached a very high performance score (*R*^*2*^ = 0.99). We next assessed the performance of the CNN for the different configurations of the multicompartment neuron network, i.e., cell morphologies, distribution of presynaptic inputs and position of the recording electrode (Table 2 B-D). Compared to the best performing linear proxy, *ERWS2*, the CNN provided an increase of performance between 2 and 8 % in most scenarios.

To gain insight into how AMPA and GABA inputs interact with layers of the network, we inspected the weights learned by different filters of the convolutional layer, as illustrated in Fig 8 for some examples of representative filters, depicted both in the time domain (panel A) and frequency domain (panel B). We observed that the majority of filters perform a band-pass and high-pass filtering of AMPA and GABA inputs and their peak frequencies are within the range [10^2^, 10^3^] Hz. This indicates that the CNN primarily uses the fast dynamics of the current inputs to construct an estimate of the EEG signal. We then asked whether we could disentangle the different transformation functions applied by the CNN to each type of input current. In signal processing, the impulse response of a linear system is typically used to understand the type of transfer function implemented by the system. Although the convolution of the first network layer is linear, subsequent network are non-linear. However, we could use a similar methodology to characterize the transformation function of the CNN by collecting the network responses to all possible combinations of unit impulses applied either to the AMPA or GABA inputs (Fig 8 C). To extract a measure of the time shift applied by the network to AMPA and GABA inputs, we computed, for each unit impulse, the difference between the time when the impulse is applied and the time in which the absolute response of the network reaches its maximum. The histogram of time shifts applied to AMPA and GABA inputs (Fig 8 D) shows that the CNN generally estimated the EEG signal by time shifting AMPA and GABA currents within the range [−2, 2] ms and the time shift could be either positive or negative.

**Fig 8.**
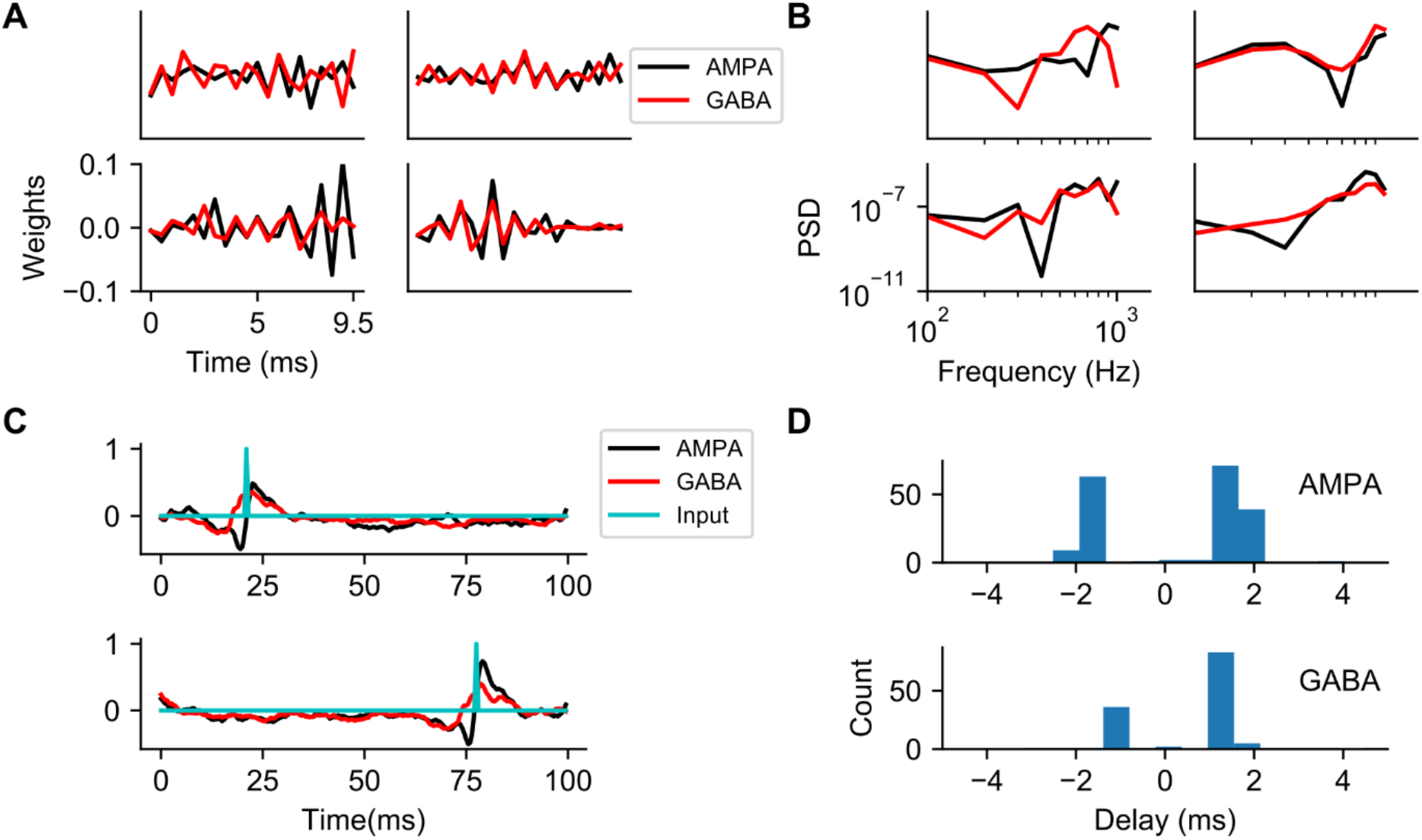
Learned filters of the convolutional layer and illustration of time shifts applied by the CNN to AMPA and GABA input currents. Examples of weights learned by four filters of the convolutional layer, depicted both in the time (A) and frequency domains (B) for the AMPA and GABA inputs. (C) Examples of the CNN outputs in response to unit impulses applied either to the AMPA or GABA inputs. (D) Histograms of time shifts applied to the AMPA and GABA inputs for all combinations of impulses. Each time shift is computed as the difference between the time when the impulse is applied and the time in which the absolute response of the CNN reaches its maximum.

### Prediction of the stimulus-evoked EEG

Evoked potentials are a useful technique that measures the transient response of the brain following presentation of a stimulus. Although the proxies we obtained have been optimized on long stretches of steady-state network activity, we investigated how well the proxies approximate an EEG evoked potential produced by a transient input. Fig 9 shows the spiking activity of the point-neuron network (panel A) and the ground-truth EEG (panel B) in response to a transient spike volley with a Gaussian rate profile applied to the thalamic input. This transient input simulates the thalamic input that reaches cortex when an external sensory stimulus is presented. A comparison of the performance obtained for all proxies is shown in panel C, while the outputs of *ERWS1*, *ERWS2* and the CNN are depicted in panel D, as an example, overlapped with the ground-truth EEG. We found that most of the current-based proxies approximated well the EEG when applying a transient burst of spikes of thalamic input, in particular ∑|*I*|, *ERWS1* and *ERWS2* which reached a performance of *R*^*2*^ = 0.9. These results suggest that these types of proxies could also be employed to predict the type of transient response seen in evoked potentials.

**Fig 9.**
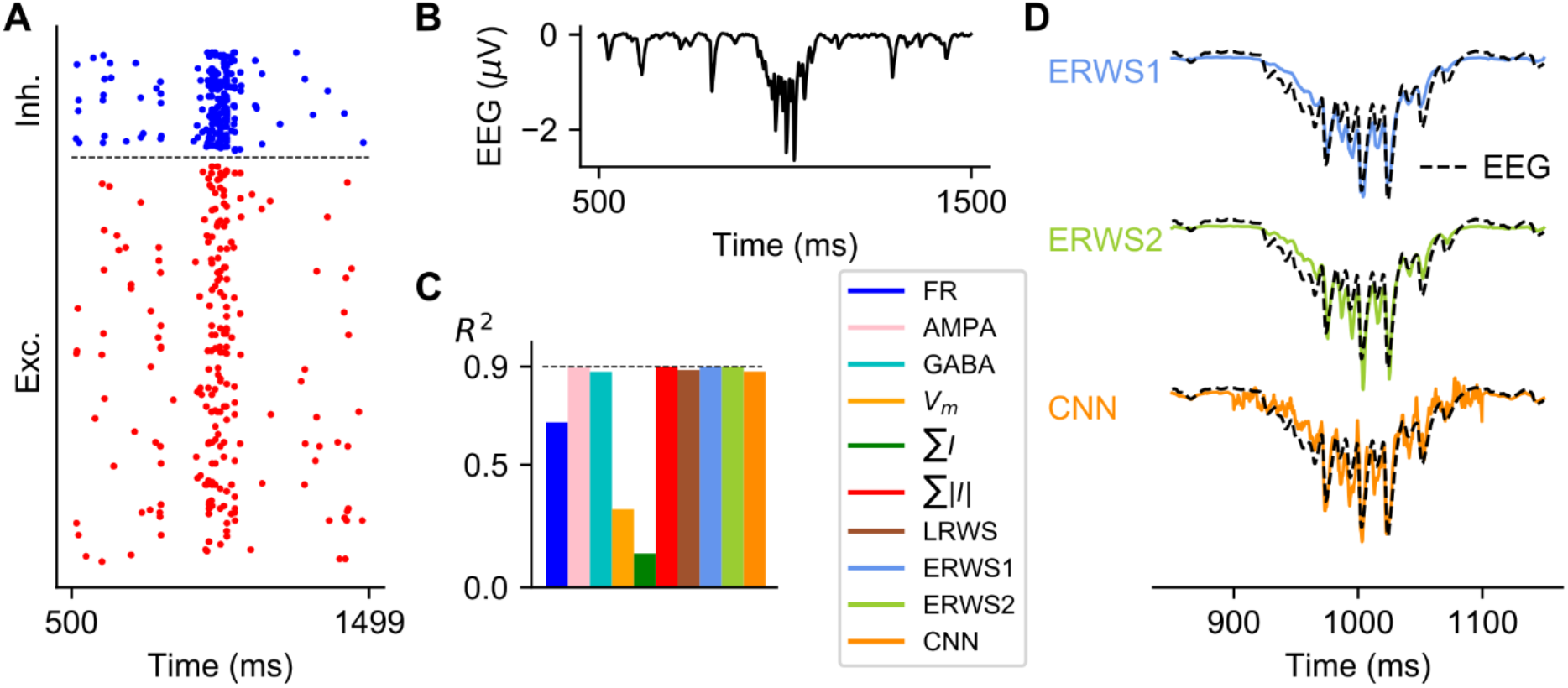
Transient activation of thalamic input with a Gaussian pulse packet. (A) Raster plot of spiking activity from a subset of cells in each population in response to a transient spike volley with a Gaussian rate profile (σ = 30 ms) centered at 1000 ms. (B) Ground-truth EEG at the top of the head model. (C) Performance of proxies calculated between 850 and 1150 ms. (D) Outputs of *ERWS1*, *ERWS2* and the CNN compared to the ground-truth EEG.

## Discussion

Understanding how to interpret experimental EEGs in terms of neural processes ultimately requires being able to compute realistic EEGs from simple and tractable neural network models, and then comparing the predictions of such models with data. Here we contributed to the first goal by developing simple yet robust and accurate methods to compute EEGs from recurrent networks of LIF point neurons, a model widely used to study cortical dynamics. We developed new linear and non-linear proxies that estimate the EEG from simple recurrent network models. A careful validation of these proxies revealed that they can give particularly accurate reconstructions of both steady-state and transient EEGs over an extensive range of network states, different morphologies, synaptic distributions and positions of the EEG electrode. These proxies thus provide a well-validated and computationally efficient way for computing a realistic EEG by simply using the output variables from simulation of point-neuron network models.

### Robustness and generality of the EEG proxies across network states, cell morphologies, synaptic distributions, and electrode locations

In many neural models used to study EEGs and LFPs, such as neural mass models [26], spiking network models [23, 29, 30] or dynamic causal models [25], extracellular potentials are simply modeled as the average firing rate or average membrane potential of excitatory neurons. While these assumptions are often reasonable, their effectiveness in describing the EEG has not been systematically validated. Here we found that these two established ways of computing the EEG worked reasonably well only under very specific conditions. However, in agreement with previous results obtained for the LFP [14, 42], we found that, for the EEG, proxies based on combinations of synaptic currents work much better and in more general conditions than proxies based on firing rates or membrane potentials. This suggests that approximations of EEGs based on firing rates or membrane potentials should be discouraged, and replaced with the use of synaptic currents, whenever possible.

Our focus has been on optimizing EEG proxies that are based on synaptic currents. The main result has been the successful development of a new class of EEG proxies, based on either an optimized linear (*ERWS1* and *ERWS2*) or non-linear (CNN) combination of time-shifted AMPA and GABA currents. We have systematically compared the performance of the new proxies in approximating the EEG with that of previous proxies used in the literature, across a range of network states, cell morphologies, synaptic distributions, and position of EEG recording electrode.

We found that, unlike all previous proxies, our new optimized EEG proxies work remarkably well for a whole range of network states which capture many patterns of oscillations, synchronization, and firing regimes observed in neocortex [12]. Predicting well the EEG over a wide range of states is important because, in many cases, EEGs are experimentally used to monitor changes in brain states, and thus models used to interpret EEGs must be able to work well over multiple states.

Our proxies were optimized using a specific pyramidal broad-dendritic-tuft morphology that generates large electric dipoles. We, however, investigated how the proxies perform when changing cell morphologies and distributions of presynaptic inputs. Our proxies showed a high performance (~80% to 95% of variance explained) across all considered scenarios, only marginally affected by changes in morphology or the distribution of GABA synapses. This suggests that our work, even though it could still be improved by using larger datasets of morphologies and synaptic distribution configurations, is already sufficiently general to accurately capture the contribution to the EEG of some major types of pyramidal neurons.

We also validated the performance of EEG proxies against changes in position of the recording electrode, with respect to the position chosen to train the proxies. The performance of proxies experienced only a moderate decrease as the position of the EEG electrode was shifted from the top of the head because of the progressive reduction in EEG amplitude. Nevertheless, the *R*^*2*^ value of *ERWS2* was maintained above 0.9 for displacements of the electrode smaller than 5 mm.

We finally demonstrated that our proxies, although trained on steady-state activity, can approximate well EEG evoked potentials, capturing the transient dynamics in response to stimuli and suggesting that our work could be relevant to model transient brain computations such as the coding of individual stimuli or attentional modulations.

Previous work [42] used a similar approach based on optimizing a linear proxy to predict the LFP. We extended this work by computing the EEG, rather than the LFP, and this implies that we used a head model that approximates the different geometries and electrical conductivities of the head, which was not necessary for the LFP proxy. Unlike the previous work, which considered only a reduced regime of network dynamics within the asynchronous or weakly synchronous states, we generated proxies trained and validated on a wider range of network states. Our EEG proxies were also validated on different pyramidal-cell morphologies reconstructed from experimental recordings, whereas the LFP proxy was validated on synthetically generated morphologies. As a result, our new optimized EEG proxies predict well the EEG over a wide range of states and different morphologies, unlike the LFP proxy, which worked well only for a low-input-rate state and a specific morphology of pyramidal cells.

In sum, our new optimized EEG proxies provide a simple way to compute EEGs from point-neuron networks that is highly accurate, stable across network states and variations of biophysical assumptions, and relatively invariant regarding position of the recording electrode.

### Applications and impact of the new EEG proxies

Our work provides a key computational tool that enables applying tractable network models to EEG data with significant implications in two main directions.

First, when studying computational models of brain function, our work allows quantitative rather than qualitative comparison of how different models match EEG data, thereby leading to better and more objective validations of different hypotheses about neural computations.

Second, our work represents a crucial step in enabling a reliable inference, from real EEG data, of how different neural circuit parameters contribute to brain functions and brain pathologies. Since the EEG conflates many circuit-level aggregate neural phenomena organized over a wide range of frequencies, it is difficult to infer from its measure the value of key neural parameters, such as for example the ratio between excitation and inhibition [1, 53]. Developing tractable neural networks that include an explicit relationship between the EEG response and neural network parameters is a way to address this issue. By fitting such models to real EEG data, estimates of neural network parameters (such as the ratio between excitation and inhibition or properties of network connectivity) can be obtained from EEG spectra or evoked potentials. This approach could be used, for example, to test the influential theories of the excitation-inhibition balance as a framework for investigating mechanisms in neuropsychiatric disorders [54, 55], to empirically measure how this balance changes between patients with autistic disorder syndrome and control subjects [53], or to individuate the neural correlates of diseases that show alterations of EEG activity [56–60]. Thus, our EEG proxies have clear relevance for connecting EEG in human experiments to cellular and network data in health and disease.

Although more work is needed to be able to interpret empirical EEGs in terms of network models, there are several facts that indicate that our proxies can potentially help in this respect. Recent attempts to infer neural parameters from EEGs or other non-invasive signals, based on network models that use less accurate proxies than the ones developed here, are nevertheless beginning to provide credible estimates of key parameters of underlying neural circuit such as excitation-inhibition ratios [53, 61], as well as accurate descriptions of cortical dynamics. For example, previous theoretical studies have modeled the LFP/EEG as the sum of absolute values of synaptic currents [14, 15, 34, 45]. This type of proxy, though simplified, was shown to be sufficient to explain quantitatively several important properties of cortical field potentials, including the relationship between sensory stimuli and the spectral coding of LFPs [14], cross-frequency and spike-field relationships [34], and LFP phase of firing information content [15]. We thus expect that the new EEG proxies can help building on these encouraging results and further improve the biological plausibility and robustness of neural parameter estimation from EEGs.

### Linear vs non-linear proxies

We optimized the EEG proxies by training either linear or non-linear EEG prediction models based on synaptic currents. In particular, the linear proxies (*ERWS1* and *ERWS2*) were based on an optimized linear combination of time-shifted AMPA and GABA currents. Alternatively, we investigated the application of a shallow CNN that could capture more complex interactions between synaptic currents to estimate the EEG. Compared to the best performing linear proxy, *ERWS2*, the non-linear EEG proxy based on a convolutional network provided a sizeable increase of performance (2 to 8 %, see Table 2) and it provided a very high performance (>85%) in all conditions. The convolutional weights that we provide (see [62] and Section “Data and Code Availability”) can be used to easily compute these non-linear EEGs proxies using similar computational power as that employed for linear proxies. However, the drawback of CNNs is that it is harder to infer direct relationships between synaptic currents and the EEG, whereas these relationships are apparent and immediate to interpret with linear proxies (see section below). However, we showed that this problem could be in part attenuated when using tools to visualize the transformation function implemented by the CNN, which allow an understanding of how synaptic currents are transformed by the non-linear proxy.

### Limitations and future work

The present network modelling scheme involves several major assumptions with respect to simplification of the multi-layered cortical column architecture, and combined use of point-neuron and multicompartment networks.

Our proxies have been extensively validated for a model with one class of pyramidal cells and are expected to be applied to models of any brain area in which the EEG is likely to be generated by one dominant population. We chose to model a single cortical layer, layer 2/3, based on previous computational work suggesting that this layer gives a large contribution to extracellular potentials [30, 35]. Although we have shown that our proxies generalize well for different L2/3 pyramidal-cell morphologies, it will be important to extend our work to quantify contributions from other cortical laminae and cell morphologies to the generation of EEGs. In this regard, it is important to note that electrical potentials in the brain tissue add linearly and the superposition of individual contributions to the EEG is in principle straightforward to compute if the amplitude of each laminar contribution is known. Thus, we could approximate the total EEG by a suitable linear combination of individual proxies computed for each population. We envisage future studies that address this issue by coupling multi-layer spiking models of cortical circuits [30, 63, 64] with multi-layer multicompartment neuron models within the hybrid modelling scheme.

The hybrid modelling approach [30] offers the advantage that we can vary parameters of the EEG-generating model, e.g., cell morphologies or synaptic distribution, without affecting the spiking dynamics. The disadvantage of this approach is, however, that the multicompartment network does not match the point-neuron network in every respect. For instance, even though the synaptic input conductances were identical in the two models, the resulting soma potentials of multicompartmental neurons were not identical to those of the point neurons because of passive dendritic filtering or the lack of a membrane-voltage reset mechanism following spike, among other effects. This inconsistency could, at least partially, be resolved by extracting the effective synaptic weight distributions from multicompartment neurons and use them in the point-neuron network in order to make the two simulation environments even more similar [65].

Calculation of EEG signals requires a head model, and here we have used the simple analytic four-sphere head model. There are however many high-resolution, anatomically detailed, and potentially personalized head models available, which for example take into account the folded cortical surface of the human brain [66–68]. Importantly, the EEG proxies developed here can be easily used in combination with such complex head models. This is because the EEG signal calculated at top of the head in the four-sphere head model, resulting from a current dipole directly below the electrode, is in fact just a scaling of the dominant component (the component aligned with the depth axis of the cortex) of the original current dipole [35]. This means that our proxies developed for the EEG signal at the top of the head in the four-sphere model, are in fact equally valid as proxies for the (normalized) dominant component of the population current dipole moment, that is, the sum of all single-cell current dipole moments (Sup. Fig 1). Such population current dipoles can be used directly in combination with existing detailed head models to calculate EEG signals, see for example [35]. Further, note that in Fig 6, we tested that the proxy for the EEG signal optimized for the top of the head worked well for other head locations.

### Insights gained from proxies about the neural contributions to the EEGs

Parameters of the linear proxies, and their variations over cortical states, allow immediate postulations about how synaptic currents combine to generate an EEG. We showed that the time shifts of *ERWS1* and *ERWS2* resulted from the optimization process have opposite signs, indicating that the EEG signal depends on both causal and non-causal components of AMPA and GABA currents. The presence of non-causal components in a proxy may appear at first counterintuitive but as previously found for the LFP [6], this reflects the recurrent nature of interactions within a cortical circuit, which makes it impossible to separate completely cause and effects and leads to both causal and non-causal dependencies.

Importantly, the analysis of the best performing proxy, *ERWS2*, whose parameters change as a function of the external input rate, revealed that the contribution of synaptic currents to the EEG dynamically varies with the cortical state. Specifically, we found that time shifts of AMPA and GABA currents, and the relative weighting between GABA and AMPA currents depend on the network state. In particular, we observed a larger weight of GABA currents for low values of the external input and the opposite effect, stronger weight of AMPA currents, as the external rate increases. This suggests that the contribution of neural activity to the EEG is a dynamic, rather than a static process, and underlies the importance of developing EEG proxies, such as those developed here to capture these variations.

## Methods

### Overview of the approach for computing the proxies and the ground-truth EEG

Our focus is on computing an accurate prediction of the EEG (denoted as “proxy” in the following) based simply on the variables available directly from the simulation of a point-neuron network model. The point-neuron network was constructed following a well-established configuration based on two populations of LIF point neurons, one excitatory and other inhibitory, with recurrent connections between populations [12], as illustrated in Fig 1 A. The network receives two types of external inputs: a thalamic synaptic input that carries the sensory information and a stimulus-unrelated input representing slow ongoing fluctuations of cortical activity.

The ground-truth EEG (referred to simply as “EEG” in the paper) with which to compare the performance of the different proxies is here computed using the hybrid modelling scheme [30, 35, 42, 43]. We created a network of unconnected multicompartment neuron models with realistic morphologies and distribute them within a cylinder of radius *r* = 0.5 mm (Fig 1 C). We focused on computing the EEG generated by neurons with somas positioned in one cortical layer so that the soma compartments of each cell are aligned in the Z-axis, 150 μm below the reference point *Z* = 8.5 mm, and homogenously distributed within the circular section of the cylinder. In our default setting, all dendrites of inhibitory cells receive GABA synapses while only those dendrites of excitatory cells below *Z* = 8.5 mm receive GABA synapses. AMPA synapses are homogenously positioned along the Z-axis in both cell types.

EEGs were generated from multicompartment neurons in combination with a forward-modelling scheme based on volume conduction theory [6]. From each multicompartment neuron simulation the current dipole moment of the cell was extracted with LFPy [39]. Next, these current dipole moments and the locations of the cells were used as input to the four-sphere head model to calculate all single-cell EEG contribution. The ground-truth EEG signal is the sum of all such single-cell EEG contributions. To approximate the different geometries and electrical conductivities of the head, we computed the EEG using the four-layered spherical head model described in [49]. In this model, the different layers represent the brain tissue, cerebrospinal fluid (CSF), skull, and scalp, with radii 9, 9.5, 10 and 10.5 mm respectively, which approximate the dimensions of a rodent head model [46]. The values of the conductivities chosen are the default values of 0.3, 1.5, 0.015 and 0.3 S/m. The EEG electrode is located on the scalp surface, at the top of the head model (Fig 1 C).

The time series of spikes of individual point neurons were mapped to synapse activation times on corresponding postsynaptic multicompartment neurons. Each multicompartment neuron was randomly assigned to a unique neuron in the point-neuron network and received the same input spikes of the equivalent point neuron. Since the multicompartment neurons were not interconnected, they were not involved in the LIF network dynamics and their only role was to transform the spiking activity of the point-neuron network into a realistic estimate of the EEG. The EEG computed from the multicompartment neuron model network was then used as benchmark ground-truth data against which we compare different candidate proxies (Fig 1 D).

### Definition and computation of the proxies that approximate the ground-truth EEG

A proxy is defined as an estimation of the EEG based on the variables available from the point neuron model over all excitatory neurons. Unless otherwise stated, we only considered the contributions of pyramidal cells to generate the EEG (in both the point-neuron and multicompartment neuron networks). The first six proxies that we tested were those used in previous literature for predicting the EEG or the LFP from point-neuron networks. These were: the average firing rate (*FR*), the average membrane potential (*V*_*m*_), the average sum of AMPA currents (*AMPA*), the average sum of GABA currents (*GABA*), the average sum of synaptic currents (∑*I*) and average sum of their absolute values (∑|*I*|). Note that ∑*I* and ∑|*I*| are defined as the sum of both AMPA and GABA currents. Because of the opposite signs assigned to the AMPA and GABA currents, ∑|*I*| is equivalent to the difference between these currents. Computation of the average *FR* was calculated with a temporal bin width of 1 ms, and then filtered with a 5-ms rectangular window to produce a smoother output of the *FR*.

For several reasons (e.g., different rise and decay time constants or different peak conductances), we expect that AMPA and GABA currents contribute differently to the EEG and that the optimal combination of both types of currents could involve different time delays between them. Following Mazzoni and colleagues [42], the new class of current-based proxies, the weighted sum of currents (*WS*), was based on a linear combination of AMPA and GABA currents, with a factor *α* describing the relative ratio between the two currents and a specific delay for each type of current (*τ*_*AMPA*_, *τ*_*GABA*_):

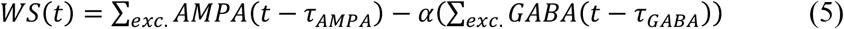

The optimal values of *α*, *τ*_*AMPA*_ and *τ*_*GABA*_ were found to be 1.65, 6 ms and 0 ms for the LFP, respectively [42]. As a result, the LFP reference weighted sum (*LRWS*) proxy was defined as

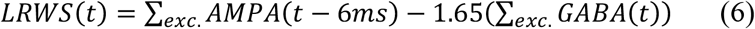

Here we also introduced two new proxies derived from the *WS* formulation: the EEG reference weighted sum 1 (*ERWS1*) and the EEG reference weighted sum 2 (*ERWS2*), whose parameters were optimized to fit the EEG under different network states of the point-neuron network. While the concept of *ERWS1* is similar to that of *LRWS*, with fixed optimal values of *α*, *τ*_*AMPA*_ and *τ*_*GABA*_, the parameters of the *ERWS2* were defined as a power function of the firing rate of the thalamic input (*v*_*0*_, unitless) to account for possible dependencies of the EEG with the external rate:

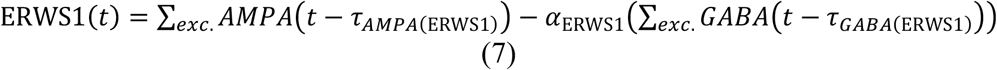

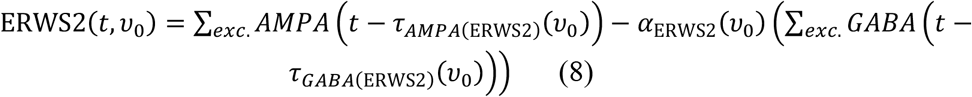

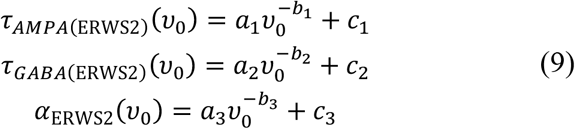

The total number of parameters to optimize was 3 for *ERWS1* (*α*_*ERWS1*_, *τ*_*AMPA*_(_*ERWS1*_) and *τ*_*GABA*_(_*ERWS1*_)) and 9 for *ERWS2* (*a*_*1*_, *b*_*1*_, *c*_*1*_, *a*_*2*_, *b*_*2*_, *c*_*2*_, *a*_*3*_, *b*_*3*_ and *c*_*3*_). We experimented with other classes of functions (e.g., exponential and polynomial functions) to describe the dependency of parameters of *ERWS2* with *v*_*0*_ but the best performance results were found with a power function.

### Leaky integrate-and-fire point-neuron network

We implemented a recurrent network model of LIF point-neurons that was based on the Brunel model [31] and the modified versions developed in subsequent publications [14, 15, 34, 42, 45, 69]. These models have demonstrated to explain well and capture a large fraction of the variance of the dynamics of neural activity in primary visual cortex during naturalistic stimulation, including a wide range of cortical oscillations such as low-frequency (1-12 Hz) and gamma (30-100 Hz) oscillations. In particular, the network structure and model parameters are the same ones used in [69] with conductance-based synapses (we refer the reader to this publication for an in-depth technical description of the implementation). Briefly, the network was composed of 5000 neurons, 4000 are excitatory (i.e., their projections onto other neurons form AMPA-like excitatory synapses) and 1000 inhibitory (i.e., their projections form GABA-like synapses). The neurons were randomly connected with a connection probability between each pair of neurons of 0.2. This means that, on average, the number of incoming excitatory and inhibitory connections onto each neuron was 800 and 200, respectively. Both populations received two different types of excitatory external input: a thalamic input intended to carry the information about the external stimuli and a stimulus-unrelated input representing slow ongoing fluctuations of activity. Spike trains of the external inputs are generated by independent Poisson processes. While the firing rate of every individual Poisson process for the thalamic input was kept constant in each simulation (within the range [1.5, 30] spikes/s), the firing rate of the cortico-cortical input was varied over time with slow dynamics, according by an Ornstein-Uhlenbeck (OU) process with zero mean:

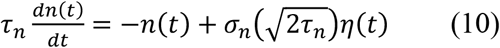

Here 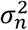 (0.16 spikes/s) is the variance of the noise, *η*(*t*) is a Gaussian white noise and *τ*_*n*_ (16 ms), the time constant. The full network description is given in Tables 3 and 4, following the guidelines indicated in [70].

**Table 3.** Description of the point-neuron network.

| A: Model summary |  |  |  |
| --- | --- | --- | --- |
| Structure | Excitatory-inhibitory (E-I) network |  |  |
| Populations | Two: excitatory and inhibitory |  |  |
| Input | 2 independent Poisson spike trains, one with a fixed rate and the other with a time-varying rate generated by an OU process |  |  |
| Measurement | Spikes, membrane potential, AMPA and GABA currents |  |  |
| Neuron model | Cortex: leaky integrate-and-fire (LIF) with fixed threshold and fixed absolute refractory time; external inputs: point process |  |  |
| Synapse model | Difference of exponential functions; conductance-based synapses |  |  |
| Topology | None |  |  |
| Connectivity | Random and sparse |  |  |
| B: Populations |  |  |  |
| Type | Elements | Size |  |
| Pyramidal cells | LIF neurons | 4000 |  |
| Interneurons | LIF neurons | 1000 |  |
| Thalamic input | Poisson generator | 1 |  |
| Cortico-cortical input | Poisson generator | 1 |  |
| C: Connectivity |  |  |  |
| Name | Source | Target | Pattern |
| AMPA<sub>Pyr_Pyr</sub> | Pyramidal | Pyramidal | Random convergent (p = 0.2), weight $g_{Pyr\_Pyr}$ |
| AMPA<sub>Pyr_Int</sub> | Pyramidal | Interneuron | Random convergent (p = 0.2), weight $g_{Pyr\_Int}$ |
| GABA<sub>Int_Pyr</sub> | Interneuron | Pyramidal | Random convergent (p = 0.2), weight $g_{Int\_Pyr}$ |
| GABA<sub>Int_Int</sub> | Interneuron | Interneuron | Random convergent (p = 0.2), weight $g_{Int\_Int}$ |
| AMPA<sub>tha_Pyr</sub> | Thalamic | Pyramidal | Fixed in-degree (800), weight $g_{tha\_Pyr}$ |
| AMPA<sub>tha_Int</sub> | Thalamic | Interneuron | Fixed in-degree (800), weight $g_{tha\_Int}$ |
| AMPA<sub>cort_Pyr</sub> | Cortical | Pyramidal | Fixed in-degree (800), weight $g_{cort\_Pyr}$ |
| AMPA<sub>cort_Int</sub> | Cortical | Interneuron | Fixed in-degree (800), weight $g_{cort\_Int}$ |
| D: Neuron model |  |  |  |
| Type | Leaky integrate-and-fire |  |  |
| Description | $\tau_m \frac{dV(t)}{dt} = -V(t) + V_{leak} - \frac{I_{tot}(t)}{g_{leak}},$ $I_{tot}(t) = \sum_{N_{AMPA_{rec}}} I_{AMPA_{rec}}(t) + \sum_{N_{GABA_{rec}}} I_{GABA_{rec}}(t) + I_{AMPA_{ext}}(t),$ | | |
| E: Synapse model |  |  |  |
| Type | Conductance-based synapse, difference of exponentials [31] |  |  |
| Description | $I_{syn}(t) = g_{syn} s_{syn}(t) (V(t) - E_{syn}),$<br>if a presynaptic spike occurs:<br>$s_{syn}(t) = \frac{\tau_m}{\tau_d - \tau_r} \left[ \exp\left(\frac{-t - \tau_l}{\tau_d}\right) - \exp\left(\frac{-t - \tau_l}{\tau_r}\right) \right]$ | | |
| F: Input |  |  |  |
| Type | Description |  |  |
| Poisson generator | Thalamic input, time-constant input with rate $\nu_0$; each neuron receives 800 independent thalamic inputs | | |
| Poisson generator | Cortico-cortical input, OU process with zero mean; each neuron receives 800 independent cortico-cortical inputs |  |  |
| G: Global simulation parameters |  |  |  |
| Simulation duration | 3000 ms |  |  |
| Temporal resolution | 0.05 ms |  |  |
| Startup transient | 500 ms |  |  |

**Table 4.**
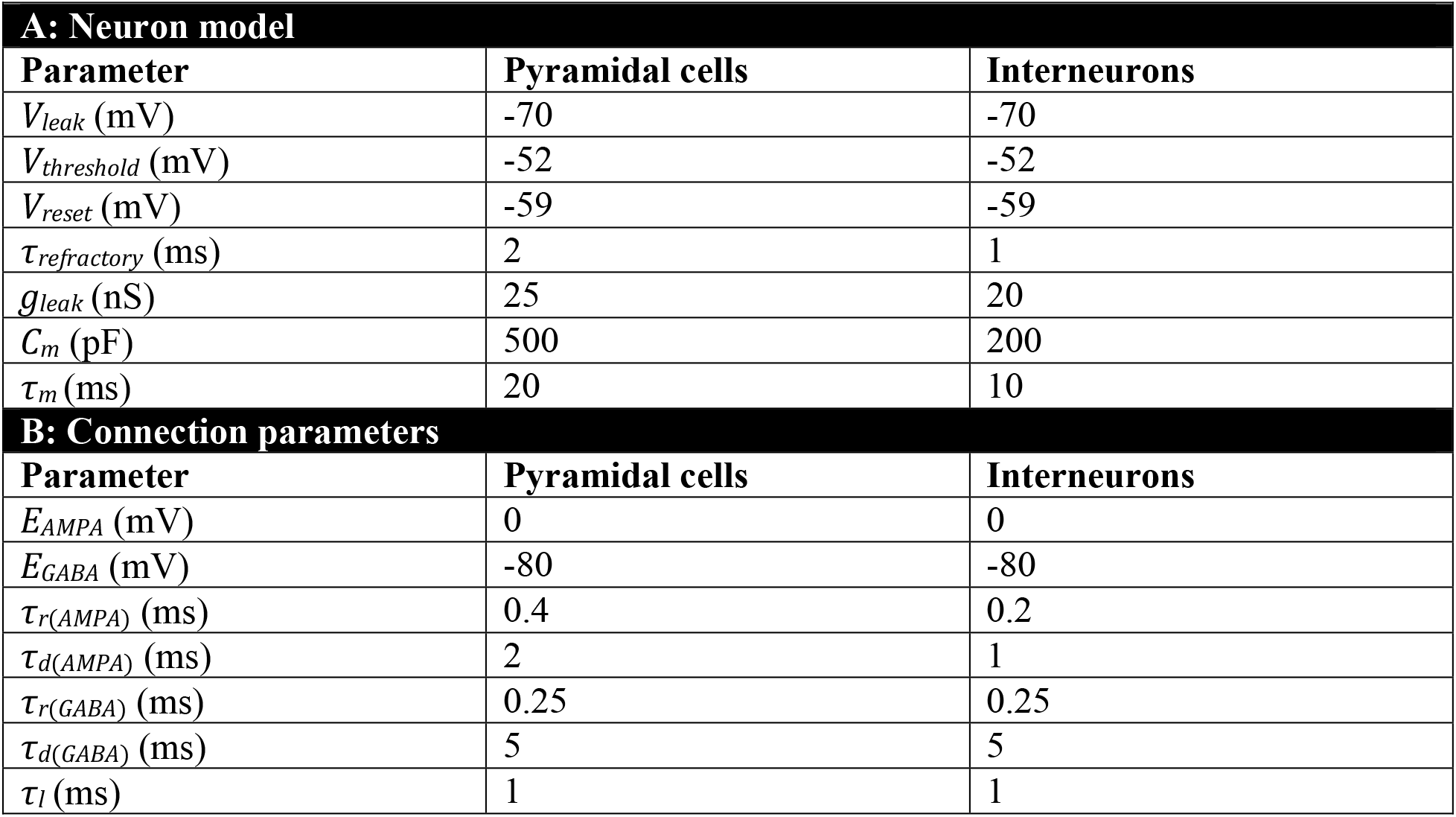
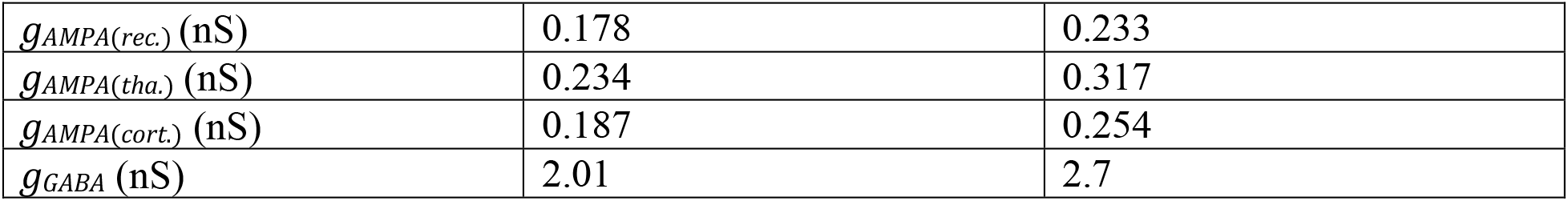
Parameters of the neuron models used in the point-neuron network.

### Multicompartment-neuron network

The EEG was computed by projecting the spiking activity of the point-neuron network onto a network of multicompartment neuron models in which every multicompartment neuron is assigned a unique corresponding point neuron. A key factor for a successful representation of the EEG is selection of proper morphologies of multicompartment neurons with detailed and realistic dendritic compartments. Our focus was on computing the EEG for cortical layer 2/3 so that we acquired representative morphological reconstructions of L2/3 pyramidal cells and interneurons from publicly available repositories: the Neocortical Microcircuitry (NMC) portal [47, 48] based predominantly on the data released by Markram and collaborators [47], and the Allen Brain Atlas (ABA) [51]. We also imposed our target animal model to be the rodent model. In our simulations, we evaluated three different types of morphologies of L2/3 pyramidal cells and one morphology of a specific type of L2/3 interneuron, the large basket cell interneuron (the most numerous class in L2/3 [47], represented as PY and LBC respectively in Table 5. Unless otherwise stated, the default morphology file used for pyramidal cells in our simulations is *dend-C250500A-P3_axon-C260897C-P2-Clone_9*.

**Table 5.** Morphologies types and file identifiers used in the multicompartment neuron network model.

| Cell type | Animal species | File identifier | Source |
| --- | --- | --- | --- |
| L2/3 PY | Rat | dend-C250500A-P3_axon-C260897C-P2-Clone_9 | NMC |
| L2/3 PY | Rat | dend-C260897C-P3_axon-C220797A-P3-Clone_0 | NMC |
| L2/3 PY | Mouse | Cux2-CreERT2, ID:486262299 | ABA |
| L2/3 LBC | Rat | C250500A-I4_Clone_0 | NMC |

Soma compartments of pyramidal cells and interneurons were randomly placed in a cylindrical section of radius 0.5 mm, at *Z* = 8.35 mm. We assumed that GABA presynaptic inputs could only be located on dendritic compartments below the reference point *Z* = 8.5 mm. AMPA synapses were homogenously distributed along the Z-axis in both cell types with random probability normalized to the membrane area of each segment. This configuration resulted in an asymmetric distribution of AMPA and GABA synapses onto pyramidal cells creating a stronger current dipole moment from these types of cells. Each multicompartment neuron was modeled as a non-spiking neuron with a passive membrane [38]. Tables 6 and 7 summarize properties of the multicompartment neuron network.

**Table 6.** Description of the multicompartment neuron network.

| A: Model summary |  |
| --- | --- |
| Structure | Unconnected populations of multicompartment neurons |
| Populations | Two: pyramidal cells and interneurons |
| Input | Presynaptic spiking activity as modeled by the point-neuron network |
| Measurement | EEG, current dipole moment |
| Neuron model | Multicompartment neuron model based on the passive cable formalism |
| **Synapse model** | Difference of exponential functions; conductance-based synapses |
| **Topology** | Cylindrical volume with radius $r = 0.5$ mm |
| **Connectivity** | None |
| **B: Populations** |  |
| **Type** | Populations of 4000 pyramidal cells and 1000 interneurons |
| **Cell positions** | Soma compartments located at $Z = 8.35$ mm and randomly distributed within the circular section of the cylinder |
| **Cell orientations** | Fixed orientation with apical dendrites oriented along the Z-axis |
| **Morphologies** | Reconstructed morphologies from the NMC and ABA (Table 5); axons removed if present |
| **C: Connectivity** |  |
| No network connectivity, synaptic inputs are generated by the point-neuron network with the same synaptic parameters (Table 4) |  |
| **D: Neuron model** |  |
| **Type** | Multicompartment reconstructed morphologies |
| **Description** | Non-spiking neurons based on the passive cable formalism (except in subsection “The performance of EEG proxies depends on the neuron morphology, distribution of synapses and the type of dendritic conductances”), with membrane capacity $c_m$, membrane resistivity $r_m$, axial resistivity $r_a$ and leak reversal potential $E_L$. |
| **E: Synapse model** |  |
| **Type** | Conductance-based synapse, difference of exponentials |
| **Description** | $I_{syn}(t) = g_{syn}s_{syn}(t)(V(t) - E_{syn}),$ $s_{syn}(t) = A \left[ \exp\left(\frac{-t-\tau_l}{\tau_d}\right) - \exp\left(\frac{-t-\tau_l}{\tau_r}\right) \right],$ where $A$ is a normalization factor to give a peak conductance $g_{syn}$ |
| **F: Input** |  |
| **Type** | Spike times of spiking neuron network (including thalamic and cortico-cortical input spikes), no recurrent input |
| **Description** | All dendrites of interneurons receive GABA synapses while only those dendrites of pyramidal cells below $Z = 8.5$ mm receive GABA synapses; AMPA synapses are homogenously positioned along the Z-axis in both cell types; synapse locations are randomly assigned onto cell compartments assuming a probability proportional to the compartment’s surface area divided by the total surface area of the cell |
| **G: Global simulation parameters** |  |
| **Simulation duration** | 3000 ms |
| **Temporal resolution** | 0.05 ms |
| **Startup transient** | 500 ms |

**Table 7.** Parameters of multicompartment neurons.

| Parameter | Pyramidal cells | Interneurons |
| --- | --- | --- |
| $c_m$ ($\mu\text{F}/\text{cm}^2$) | 1 | 1 |
| $r_m$ ($\text{k}\Omega\text{cm}^2$) | 30 | 20 |
| $r_a$ ($\Omega\text{cm}$) | 100 | 100 |
| $E_L$ (mV) | -70 | -70 |

### Optimization and validation of EEG proxies

We created two different simulated datasets, one for optimization of the *ERWS1’s* and *ERWS2’s* parameters (Eqs. 7–9), and the other dataset for validation of performance of all proxies. The datasets were generated by varying the two parameters of the point-neuron network commonly used for exploration of different network states [31, 44]: the rate of the external input, *v*_*0*_, and the relative strength of inhibitory synapses, defined here as *g* = *g*_*Int_Pyr*_/*g*_*Pyr_Pyr*_. We selected 58 values of *v*_*0*_ within the range [1.5, 30] spikes/s and 3 values of *g* (5.65, 8.5 and 11.3), which encompass the different network states: asynchronous irregular, synchronous irregular and synchronous regular [12]. For every pair (*v*_*0*_, *g*), we generated three simulations of the point-neuron and multicompartment-neuron networks with different random initial conditions (e.g., recurrent connections of the point-neuron network or soma positions of multicompartment neurons). The simulated outputs from two of these network instantiations were used for the optimization dataset and the other one for the validation dataset.

Prior to comparing the EEG traces with the point-neuron model predictions, we z-scored the proxies and the EEG signal by subtracting their mean value and dividing by the standard deviation. The best parameters of *ERWS1* and *ERWS2* were calculated by minimization of the sum of the square errors *SSE* between the ground-truth EEG and the proxy for all network instantiations *i* of the optimization dataset:

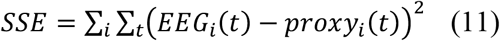

Time constants of proxies (Eqs. 7–9) were restricted to be discrete variables as the simulation time is a discrete variable. This turns the optimization problem into a discrete optimization problem, which is harder to solve than a continuous optimization problem. However, the limited number of parameters that need to be optimized allowed us to run a simple brute-force parameter search.

The performance of each proxy was evaluated by using the coefficient of determination *R*^*2*^, which is the fraction of the EEG variance explained by the proxy. *R*^*2*^ is computed as the squared value of the correlation coefficient. The validation results were calculated based on the average *R*^*2*^ of every proxy across all network instantiations *i* of the validation dataset.

### Implementation of the convolutional neural network

The processing pipeline of the CNN architecture, illustrated in Fig 7 A, was based on the machine-learning library Keras running on top of TensorFlow [71]. The CNN consists of a one-dimensional (1D) convolutional layer with 50 filters and a kernel of size 20, followed by a max pooling layer of pool size 2, a flatten layer and two fully connected layers of 200 units each (one of them is the output layer). The rectified linear unit (ReLU) function was used as the activation function for all layers, except for the output layer. To reduce overfitting, we applied L2 activity regularization (λ = 0.001) to the convolutional layer. The amount by which filters shift, the strides, is set to 1 for the convolutional layer and 2 for the max pooling layer. The input layer was formed by two channels of 1D data that correspond to the AMPA and GABA time series simulated by the point-neuron network. Instead of using data of the whole simulation (3000 ms), we split time series into multiple chunks (i.e., samples) of 100 ms, a window size that we found convenient to improve estimation accuracy of the CNN. Nodes of the output layer predict segments of the EEG signal at each 100-ms window.

The CNN was trained by first-order gradient descent (Adam optimizer [52]) with default parameters as those provided in the original paper. We defined the loss function for training as the mean squared error (MSE) between the predicted and the true values of the EEG. To monitor training, we employed the MSE and also the mean absolute error (MAE) and the coefficient of determination, *R*^*2*^. The CNN is trained for a sufficiently large number of epochs, 100 epochs, to ensure convergence of the error metrics. To train and test the CNN, we use the same datasets generated for optimizing parameters of the current-based proxies, as described above.

### Analysis of network states

To characterize the different network states of activity in the point-neuron network at the level of both single neurons and populations, we employed the descriptors developed by Kumar and collaborators for conductance-based point-neuron networks [44].

#### Synchrony

We quantified the synchrony of the population activity in the network as the average pairwise spike-train correlation from a randomly selected subpopulation of 1000 excitatory neurons. The spike trains were binned in non-overlapping time windows of 2 ms.

#### Irregularity

Irregularity of individual spike trains was measured by the coefficient of variation (the ratio of the biased standard deviation to the mean) of the corresponding interspike interval (ISI) distribution. Low values indicate regular spiking; a value of 1 reflects Poisson-type behavior. The irregularity index was computed for all excitatory neurons.

#### Mean firing rate

The mean firing rate was estimated by averaging the firing of all excitatory cells, and was calculated with a bin width of 1 ms.

### Post-processing and spectral analysis

The z-scored EEG signals and proxies are resampled by applying a fourth-order Chebyshev type I low-pass filter with critical frequency *f*_*c*_ = 800 Hz and 0.05 dB ripple in the passband using a forward-backward linear filter operation and then selecting every 10th time sample. The estimate of the normalized power spectral density (normalized PSD) was computed using the Fast Fourier Transform with the Welch’s method, dividing the EEG z-scored data into eight overlapping segments with 50 % overlap.

### Numerical implementation

Here we summarize the details of the software and hardware used to generate the results presented in this study. Point-neuron network simulations were implemented using NEST v2.16.0 [72]. EEG signals were computed using LFPy v2.0 [39] and simulations of multicompartment model neurons using NEURON v7.6.5 [73]. The CNN is constructed based on the machine-learning library Keras v2.3. The source-code structure relies on the freely available, object-oriented programming language Python (v2.7.12). Every simulation was parallelized using either a 60-CPU 256-GB server at the Istituto Italiano di Tecnologia (IIT) or the Stallo high-performance computing facilities (NOTUR, the Norwegian Metacenter for Computational Science). Simulations of the point-neuron network were performed based on thread parallelism implemented with the OpenMP library. Network simulations with NEURON used distributed computing built on the MPI interface. Computation time for completing simulations of both network models and the post-processing of results was 2 hours on average for each experimental condition. The source code to reproduce these results will be made publicly available upon final publication of this manuscript [62].

## Acknowledgments

We would like to thank M. Libera for his technical support.

## Funding

This project has received funding from the European Union’s Horizon 2020 research and innovation programme under the Marie Skłodowska-Curie grant agreement No 893825, the NIH Brain Initiative (grants U19NS107464 and NS108410), the Simons Foundation (SFARI Explorer 602849), the European Union Horizon 2020 Research and Innovation Programme under Grant Agreement No. 785907 and No. 945539 [Human Brain Project (HBP) SGA2 and SGA3], and the Norwegian Research Council (NFR) through NOTUR - NN4661K.

## Data and Code availability

The code used to generate the simulations and to perform the analyses, as well as the weights of the optimized convolutional neural networks for the EEG are available from GitHub. Martínez-Cañada, P. Github source-code repository (2020). Available from: https://github.com/pablomc88/EEG_proxy_from_network_point_neurons.

## Author contributions

Conceived project: P.M.C., S.P. Developed Methodology: all authors. Software implementation and data analysis: P.M.C. Wrote the paper (original draft): P.M.C., S.P. Wrote the paper (review and editing): all authors. Supervised project: S.P., T.F., G.T.E. Funding acquisition: S.P., T.F., P. M.C, G.T.E.

## Supplementary figures

**Supplementary Figure 1.**
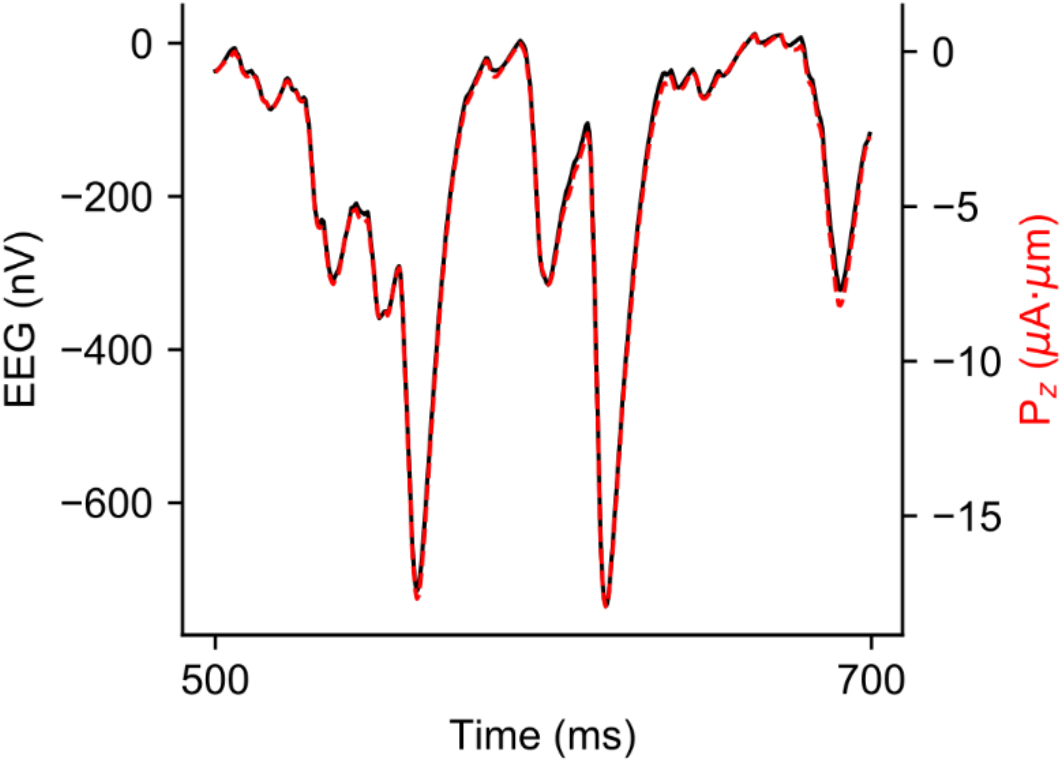
EEG (black line) and z-component (red dashed line) of the current dipole moment (P_z_) calculated at the top of head model.

## Notes

https://github.com/pablomc88/EEG_proxy_from_network_point_neurons

### Competing Interest Statement

The authors have declared no competing interest.

